# Identifying rare spontaneous mutations through wildtype *E. coli* population sequencing

**DOI:** 10.1101/2025.07.08.663732

**Authors:** R Green, M Bawn, A Angus-Whiteoak, M Jago, FJ Whelan, M Lagator, N Hall, CG Knight, R Krašovec

## Abstract

Understanding the rate and types of mutations occurring in different populations is fundamental to our ability to predict and potentially manipulate evolution. Common techniques for microbial mutation rate estimation fall into 2 camps; rapid, locus specific, generally liquid-culture based fluctuation assays; and slow, genome-wide, generally solid-culture based mutation accumulation experiments. This constrains the hypotheses which can be tested, notably the mutagenic effects of liquid environments cannot be rapidly quantified at a genome-wide scale. One example of such effect is the negative association previously observed between population density and mutation rates at many marker loci using the fluctuation assay. Because homogeneous population density is specifically relevant in liquid culture, this association has not been tested at a truly genome-wide scale. We fill this methodological gap by developing a novel pipeline that relies on population sequencing to capture how mutation rates are affected by population density in liquid cultures. We simulate expected mutation counts in a growing population along with random sampling during sequencing to estimate the necessary sequencing coverage as ≥1,000-fold. We then carry out PCR-free sequencing of 95 wildtype *E. coli* populations at this coverage, calling rare variants with both reference-based and reference-free methods. These variant-calling methods are prone to different sources of error which can be minimised by considering only mutations called by both pipelines. This approach identifies 119 mutations across all cultures, with (non)coding/(non)synonymous and mutational spectrum profiles consistent with being samples of spontaneous mutation unbiased by selection. The distribution of these mutations also supports the motivating hypothesis, finding that mutation counts across the genome are strongly negatively associated with population density. This demonstrates the utility of population sequencing for the rapid testing of many previously inaccessible evolutionary biology hypotheses.

## Introduction

### The importance of estimating mutation rates

Mutations provide the fundamental fuel of evolution; it is therefore a key evolutionary question to estimate the rate at which mutations occur (**μ**) and how this varies under different environmental conditions. *De novo* mutations can provide access to important traits such as antibiotic resistance (e.g. Garibyan et al. 2003) making their study directly relevant to improving health outcomes. While recent progress has been made in using population sequencing to identify non-fixed mutations in order to estimate mutation rates in cancer (Caravagna et al. 2020) and viruses (Furuyama et al. 2025), most bacterial research in this field has yet to effectively leverage modern sequencing technology (though see for example Fusco et al. (2016) and Jee et al. (2016)). This limits research to those hypotheses which can be tested using traditional methods. Specifically, there is a trade off between methods which rapidly measure mutation rates at a single target locus, generally in liquid media, versus methods which take multiple months to fix mutations genome-wide, generally on solid media. Therefore hypothesis tests which require rapid genome-wide mutation rate estimates in liquid culture are not easily accessible.

We propose that specific features of both existing methods can be combined by using population sequencing of liquid cultures after a single growth cycle to provide rapid, genome wide assessment of mutational patterns. All methods have to face biases introduced by natural selection and drift changing the frequency of any mutation as soon as it’s arisen, obscuring the rate of the underlying mutational process. Our approach also overcomes biases introduced by natural selection by limiting the time allowed for growth and considering mutations across the whole genome, not just at a phenotypically observable loci where fitness effects are more likely than across the genome at large (Robert et al. 2018; Soley et al. 2023).

### Overview of existing methods

Two standard methods for measuring microbial mutation rates are the **<u>fluctuation assay</u>** (FA) (Luria and Delbrück 1943) and **<u>mutation accumulation</u>** (MA) (Muller 1928; Lynch et al. 2008) experiments (reviewed by Foster 2006) (Figure 1). In almost 100 years since their conception, mutation accumulation experiments have provided important insights including the magnitude of genetic load (Kibota and Lynch 1996; Trindade et al. 2010), the mechanisms determining mutation rates (Lee et al. 2012; Foster et al. 2015) and the distribution of fitness effects of mutations under differing mutational biases (Sane et al. 2023). Similarly, fluctutation assays have enabled the discovery of many important phenomena including plasmid-induced mutagenesis (Li et al. 2019), the evolutionary dynamics of mutation rates (Sniegowski et al. 1997; Swings et al. 2017) and the impact of environmental conditions on mutation rates (Maharjan and Ferenci 2018). Despite their obvious utility, both methods have substantial drawbacks.

**Figure 1:**
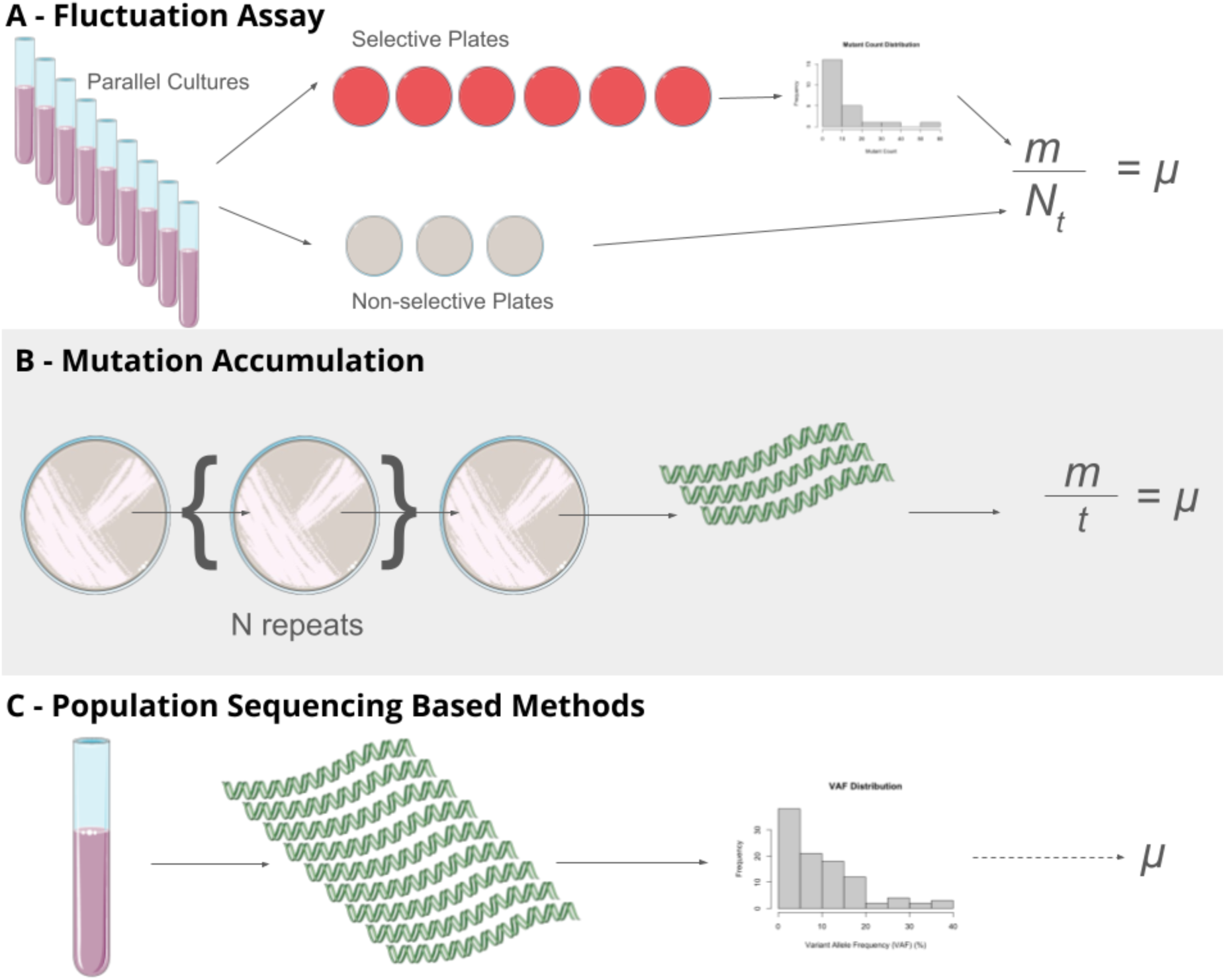
Schematic of methods for measuring microbial mutation rates. In the FA, *m* is the estimated expected number of mutational events occurring in each parallel culture, *N_t_* is the final total population size of each parallel culture and *μ* is the mutation rate. In the MA, *m* is the number of mutations and *t* is the number of generations between initiation of the mutation accumulation experiment and sequencing of descendant clone(s). The distribution of mutant counts in the top panel and the variant allele frequencies (VAF) of observed mutations in the bottom panel are simulated mock data. Green strands in the middle and bottom panels represent DNA for sequencing with depth required qualitatively indicated by the number of strands. Images sourced from bioicons.com are: testtube-purple, dna-7, petri-dish-top-red, petri-dish-top-gray and petri-dish-with-bacteria-gray icons by Servier https://smart.servier.com/ and are licensed under CC-BY 3.0 Unported https://creativecommons.org/licenses/by/3.0/.

We will now outline the standard MA and FA approaches to illustrate their respective advantages and disadvantages as summarised in Table 1.

**Table 1:** Key features of MA, FA and deep sequencing experiments based on *E. coli*.

| Feature | Fluctuation Assay (FA) | Mutation Accumulation (MA) | Population Sequencing |
| --- | --- | --- | --- |
| Scale | Single locus | Genome-wide | Genome-wide |
| **Run time** | 1 growth cycle | 30+ growth cycles | 1 growth cycle |
| **Impact of selection** | Fitness effects of chosen locus | Selection during colony growth and beneficial allele surfing | Selection during liquid growth |
| **Usual Substrate** | Liquid media | Solid media | Liquid media |
| **Key biases** | Mutation rates and spectra are only measure at a single locus which may not be representative of the genome as a whole | Depletion of deleterious alleles through natural selection. Selection acts more effectively in spatially structured colonies which are the usual growth mode in MA | False positive variant calls through A) sequencing error and B) misalignment of homologous regions |
| **Measure ment of mutation al spectrum as standard** | No | Yes | Yes |

### Pros and Cons of the Fluctuation Assay

A fluctuation assay allows the experimenter to measure the mutation rate at a single genetic locus. This measurement is generally taken as a proxy of the genome-wide mutation rate and various loci have been demonstrated to alter mutation rates under both MA and FA approaches, validating the use of single loci in this way (Sniegowski et al. 1997; Lee et al. 2012; Foster et al. 2015; Krašovec et al. 2017). The locus must have a visible phenotype, usually allowing survival under certain conditions e.g. antibiotic resistance or the ability to grow on a previously unpermissive nutrient source (Maharjan and Ferenci 2018). A single fluctuation assay involves growing multiple parallel cultures before plating the entire population of each culture on a previously lethal environment. Only a single growth cycle is required with experiments using *E. coli* generally taking ≤5 days to complete (Krašovec et al. 2019). From the distribution of observed mutant survivors on these plates the average number of mutations providing the survivor phenotype per culture can be co-estimated with the fitness effect of this phenotype (Hamon and Ycart 2012). However, the co-estimation of mutation rates and fitness effects can be unstable and other variables such the effect of death during the growth cycle must also be accounted for (Hamon and Ycart 2012). This method for rapid comparisons of mutation rates between treatment groups however, if an estimate of the per base pair mutation rate is required then the target size of mutations to the chosen phenotype must also be known and accounted for. Although FA experiments do not provide information on the mutational spectrum by default this can be achieved by selecting a phenotype which can only be accessed by a certain mutational class (Pauly et al. 2017) or by amplifying and sequencing the target site of 1 mutant per parallel culture for phenotypes accessible by multiple mutational classes (Garibyan et al. 2003; Maharjan and Ferenci 2018; Gifford et al. 2024).

### Pros and Cons of Mutation Accumulation Experiments

In contrast to FA experiments, MA allows the experimenter to identify mutations across the whole genome, with no requirement for a phenotypic marker. An MA experiment measures mutation rates by serially bottle-necking a population down to a single individual over many generations to minimise selection while mutations are allowed to accumulate. These mutations can then be identified by whole genome sequencing of individual clones after many passages to identify fixed mutations. The mutation rate per generation can then be estimated by dividing the number of fixed mutations by the total number of generations. Because mutations are identified by genome sequencing, the spectrum of mutations (i.e. the relative rates of different mutational classes including transitions, transversions and indels) is also measured. The long time-period required by such experiments is a result of low mutation rates per division. Experiments with wildtype *E. coli* often last for multiple months e.g. 100-200 days (Lee et al. 2012), 150-440 days (Foster et al. 2015) and 39-300 days (Sane et al. 2023).

Significant selection occurs during the growth of a visible colony between each bottleneck, with spatial structure within colonies grown on solid media increasing the effect of selection compared to well-mixed populations (Fusco et al. 2016; Mahilkar et al. 2022; Schreck et al. 2023). Although MA corrections to account for selective bias in well-mixed populations have been proposed, these do not account for the amplifying effect of spatial structure (Wahl and Agashe 2022). Methods for carrying out mutation accumulation in liquid culture have recently been demonstrated by (Baehr et al. 2025) marking the desire for genome-wide, liquid based mutation rate methods. The authors show consistent results between liquid and, traditional, solid MA experiments, suggesting the effect selection not to be so enhanced on solid media as previously thought. However, the constraint of long experiment run times remains and, in the case of (Baehr et al. 2025), is only overcome by using a mismatch repair deficient mutator strain.

### Population Sequencing as an Alternative Approach

It may be possible to combine the short experimental times of a fluctuation assay with the genome-wide scale of mutation accumulation experiments. Following growth from a small inoculum a population in 1mL of rich media will be significantly heterogeneous as ∼1×10^9^ genome replications will have occurred, each with the chance for mutations to occur. This naturally heterogeneous population can then be sequenced to identify rare mutations, allowing the rate and spectrum of mutagenesis to be estimated at a genome-wide scale. This experiment should only take 3 days plus sequencing time and is essentially an FA experiment where the target site is the whole genome.

We test this approach using the motivating example of density associated mutation rate plasticity. This trait is only relevant in liquid culture, and therefore cannot be observed at a genome-wide scale using traditional methods (Table 1). Using mutation rate estimates from mutation accumulation experiments, the current gold standard, we simulate the expected number of observed mutants across a variety of sequencing coverages. These calculations indicate that coverage of ≥1,000-fold is necessary for significant numbers of mutations to be observed. We then carry out population sequencing at around 1,000-fold coverage for 95 wildtype *E. coli* populations, varying population density through altering nutrient provisions. Mutations in these populations are then called using one reference-based and one reference-free variant calling pipeline. Identifying rare mutations is inherently prone to false-positive calls however, we find that these contrasting pipelines are prone to different sources of error and, therefore, by considering only mutations called by both methods signatures of bias can be reduced. Finally, we estimate the association between cell density and observed mutations per sample, demonstrating a significant negative association as observed in single locus fluctuation assays.

## Results

### Simulations of Show Coverage of ≥1000 Fold to be Necessary for Mutant Observation

The mutation rate of wildtype *E. coli* is estimated to be ∼1×10^-3^ per genome per division by mutation accumulation experiments (Lee et al. 2012). Using this estimate we can, given the initial population size (N_0_), calculate the expected number of mutations per generation for any growing *E. coli* population. As an illustrative example, we will consider a starting population of 1000 and 4 generations of growth resulting in a final population of 16,000 cells (Table 2).

**Table 2:** Example of mutations occurring over 4 generations of population growth. Here the final population size (N_t_) will be 16,000 (1,000 x 2^4^). During the growth of this population 1+2+4+8 = 15 mutations (column 3) will have occurred across the 15,000 divisions (N_t_ - N_0_). This gives us a per genome mutation rate of 1×10^-3^ (15/15,000) as expected. However, because of mutant growth 8 x 4 = 32 cells will be mutants at the end (column 6). Column 6 is invariable as earlier generations will see less mutations but more time for those mutant clones to expand, these factors cancel out result in 8 mutant cells derived from each generation in the final population.

| Generation (1) | Number of Divisions (2) | Expected number of mutations (3) | Number of remaining generations (4) | Number of final copies per mutant clone (5) | Number of mutant cells from this generation at the end (6) |
| --- | --- | --- | --- | --- | --- |
| 1 | 1000 | $1 \times 10^{-3} \times 1000$<br>= 1 | 3 | $2^3=8$ | $1 \times 8=8$ |
| 2 | 2000 | $1 \times 10^{-3} \times 2000$<br>= 2 | 2 | $2^2=4$ | $2 \times 4=8$ |
| 3 | 4000 | $1 \times 10^{-3} \times 4000$<br>= 4 | 1 | $2^1=2$ | $3 \times 2=8$ |
| 4 | 8000 | $1 \times 10^{-3} \times 8000$<br>= 8 | 0 | $2^0=1$ | $4 \times 1=8$ |

Given the information in Table 2 we can estimate the number of variants which will be sequenced at X depth for any given sequencing coverage. For each generation we will use a binomial distribution to estimate the probability of X reads being observed for any individual clone given the coverage and the mutant frequency (probability of success). The probability of an individual clone being observed at X depth can then be multiplied by the expected number of mutations in the given generation to give the total expected number of clones at X depth from this generation. This estimate of observed clones at depth X can then be summed across all generations for a population wide estimate of the number of observed clones at depth X. In (Figure 2) we take an example population of 1000 initial cells growing for 20 generations to a final population size of ∼1×10^9^ and estimate the number of clones at depths of 1 to 10 for sequencing coverage of 100 to 1×10^5^ for both wildtype *E. coli* and a mutator strain with a 100-fold increased mutation rate. Note that for clarity we will refer to the total number of reads mapping to a given position in the genome (reads per basepair) as coverage and the number of reads carrying a specific variant allele as depth. For example, if there are 100 reads mapping to a single locus and 20 carry a variant allele then we will say that the coverage is 100 fold and the depth of the variant allele is 20.

**Figure 2:**
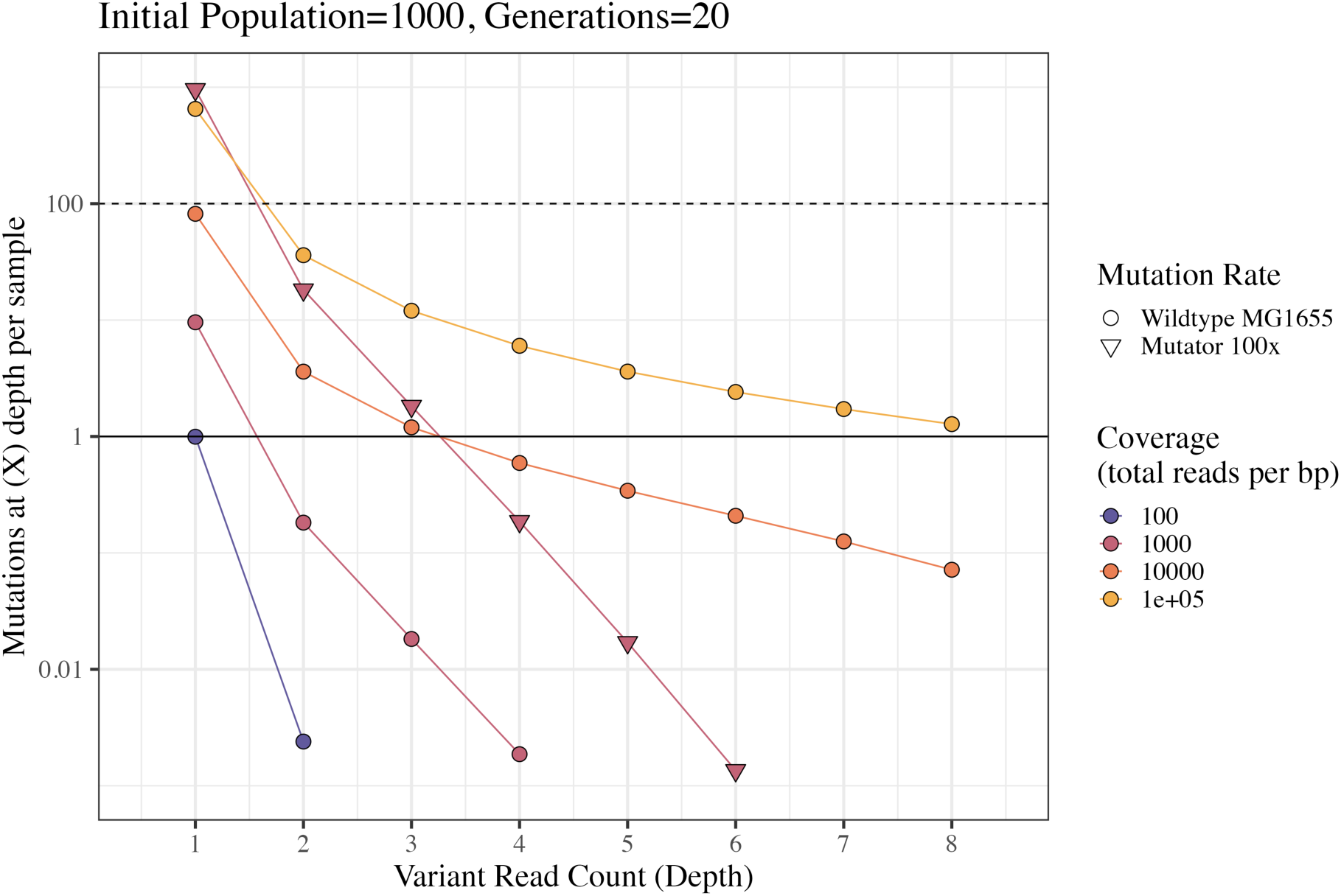
Predicted number of mutations observed at variant depth of 1 to 8 from a population initiated by 1000 cells and grown for 20 generations. Sequencing coverage of 100, 1000, 10000 and 100000 are shown in blue, maroon, orange and yellow respectively. Estimates based on the wildtype mutation rate of *E. coli* MG1655 (1×10^-3^ per genome per division) are shown by circles. Estimates for a mutator with a 100-fold higher mutation rate are shown for sequencing coverage of 1000 only as triangles. R code for all simulated data shown here can be found in supplementary data file manuscript.qmd. Note log scale y-axis.

From Figure 2 we can see that greater sequencing coverage will increase the number of mutant clones identified. While sequencing coverage of 100 is not able to identify ≥1 mutation(s) at any variant depth, coverage of 1×10^5^ is expected to identify ≥1 mutation(s) per sample at all depths up to 8. Depending on the stringency of variant calling, mutations with particularly low depth may be discarded as being more likely to derive from sequencing errors. Standard bacterial whole genome sequencing generally aims for coverage of 10-50 fold however, 1,000 fold coverage is relatively accessible target with modern technology. Using 1,000 fold coverage we see that 9.6 mutant clones are expected to be identified at a depth of 1 while only 0.18 clones are expected at a coverage of 2. In other words, 1 in 5.5 samples should carry a mutation at a depth of 2 when coverage is 1000. In contrast, a mutator strain with a 100-fold mutation rate increase is expected to produce 18 mutations at a depth of 2 given 1000 fold coverage. This clear advantage of mutator strains is the reason for their use in existing work measuring microbial mutation rates by population sequencing (Fusco et al. 2016) and liquid mutation accumulation experiments (Baehr et al. 2025). However, mutator strains are often biased not only in their mutation rates but also in their mutational spectra (the relative rates of different mutational classes e.g. transitions vs transversions) (Foster et al. 2015; Garushyants et al. 2024). Mutator strains also frequently show different responses of mutation rate to environmental conditions to their wildtype counterparts. For example, the mutator strain **Δ***mutT* differs in its mutational response to both anaerobiosis and population density from wildtype *E. coli* (Fowler et al. 1994; Krašovec et al. 2017). Given the changes in mutational spectra and environmental responsiveness seen in mutator strains we have chosen to carry out the experiment presented here using wildtype *E. coli* MG1655.

### Rare Variants Can be Cross-Validated Using Reference-Based and Reference-Free Variant Calling Methods

To test the reliability of population sequencing based methods for measuring mutational patterns across environmental conditions we sequenced 23-24 end point populations grown in each of 4 nutrient conditions (Davis Minimal media supplemented with 100, 125, 250 or 1000 mg glucose per L). This variation in nutrient provisioning will create variation in final population density, allowing for the first genome-wide test of density associated mutation rate plasticity (DAMP). DAMP has been previously observed as a consistent negative association between population density and mutation rates at a handful of marker loci including *rpoB* (Krašovec et al. 2014), *gyrA/B* (Krašovec et al. 2017) and *cycA* (Krašovec et al. 2019). Conditions closely matched to the simulations shown in Figure 2, with initial populations of 1000 - 6500 colony forming units (CFU) grown for 18 - 23 generations. DNA from all samples was prepared using PCR-free methods prior to Illumina Novaseq sequencing (see Methods for details).

There are fundamentally two distinct ways to call variants in sequencing data: by identifying differences between reads and a known reference genome or by identifying differences between the reads as compared to one another. These approaches are likely to differ in their biases and we therefore chose to implement a representative of each approach to explore these differences. Lofreq (Wilm et al. 2012) was used as our reference-based tool and DiscoSnp++ (Peterlongo et al. 2017) as our reference-free tool. We then compared results from these contrasting approaches to identify mutations which are cross-validated by being called under both methods. Pipelines used for both the reference-based and reference-free methods are shown in Figure 3. Note that mutations called by the reference-free approach implemented by DiscoSnp++ (Peterlongo et al. 2017) are subsequently mapped back to the reference to identify the position and directionality of each mutation (e.g. identifying an A>G vs G>A mutation).

**Figure 3:**
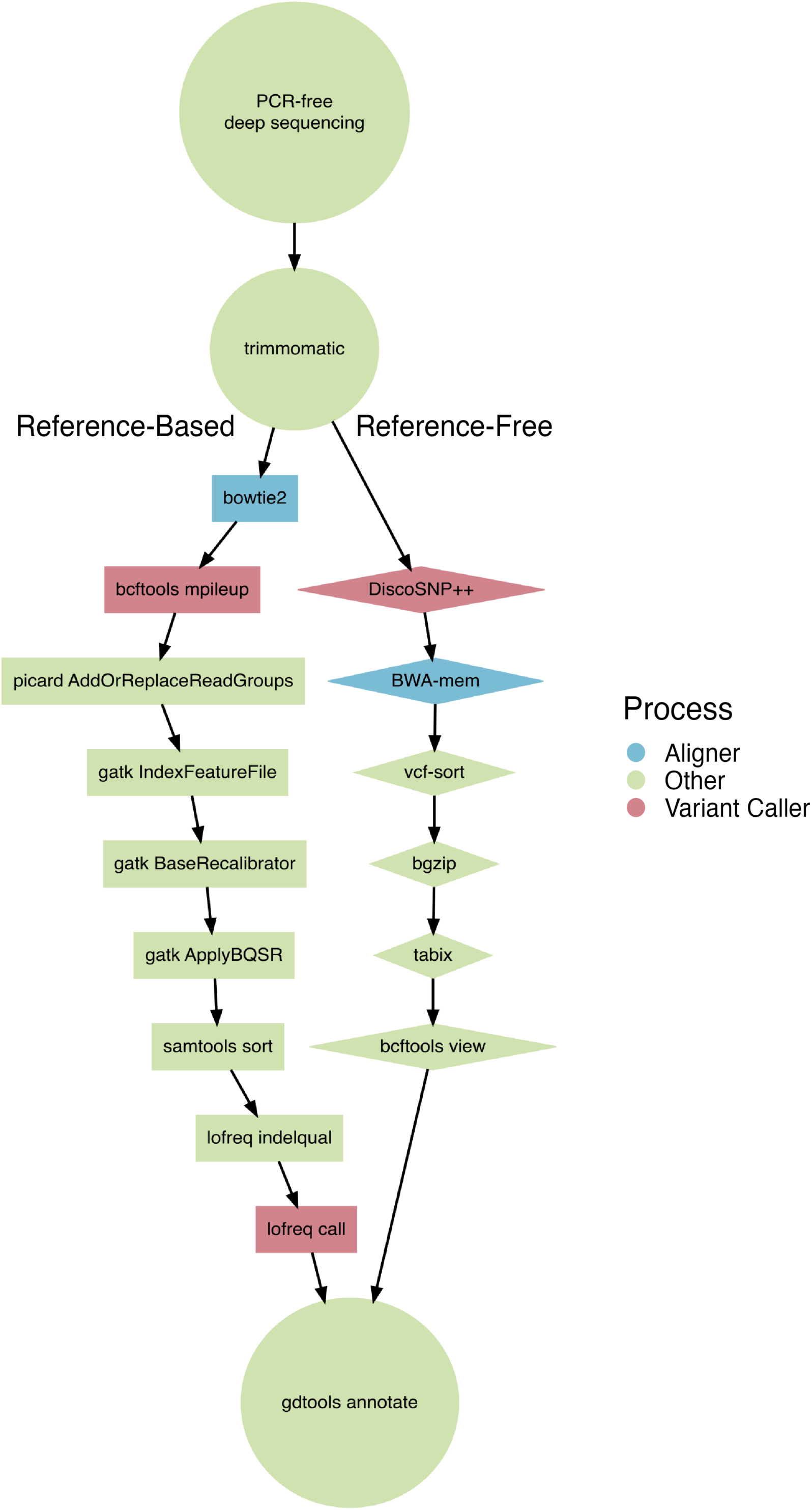
Flow diagram of reference based (left) and reference free (right) variant calling pipelines. Rectangles show steps in the reference based pipeline (based on (Jago et al. 2025)) while diamonds show the reference free pipeline using DiscoSNP++ (Peterlongo et al. 2017). Alignment steps are shown in blue, variant calling in red and all other steps in green. Full details of the options used for each tool can be found in the Methods.

We tested both the reference-based and reference-free pipelines with simulated Novaseq reads generated by InSilicoSeq (Lelieveld et al. 2024). We find that while DiscoSnp++ is more prone to calling sequencing errors as mutations it will not erroneously call any mutations if the reads given are error-free (Figure S6). In contrast while lofreq calls less errors as mutations it continues to call false positives even with error-free simulated reads (Figure S6). This difference in the false-positives to which each method is prone makes their use for cross-validating one another likely to be successful.

After calling mutations for all 95 wildtype *E. coli* samples using these two methods we find 119 cross-validated mutations called by both methods (Figure 4). We note that not only are these 119 mutations called by both methods but the variant allele depth is also very consistent across the two methods (Figure S1).

**Figure 4:**
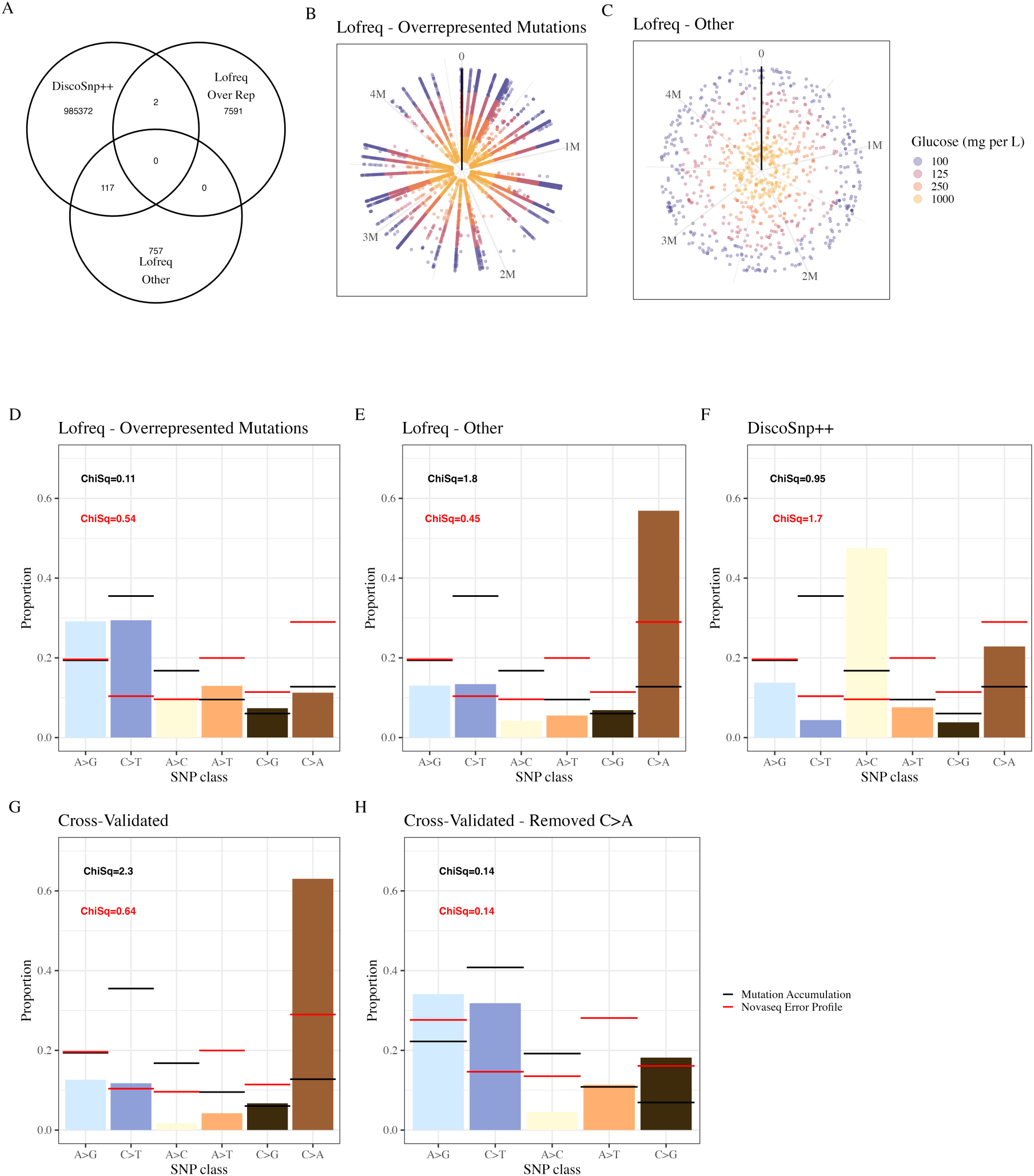
Variants called and cross validated by reference based and reference free methods. A) Venn-diagram of unique mutation counts (unique ID = sample + mutation + position) called by discosnp++, lofreq over rep (mutations called in >1 sample) and lofreq other (mutations called in only 1 sample) showing 117+2 cross-validated mutations. B and C) each concentric circle shows mutations in a single sample with radius lines formed by mutations which are called in multiple samples. D to G) For each group of called mutations the proportion of SNP mutations in each class are shown by bars with the mean wildtype mutation spectrum seen in MA Long et al. (2016), Lee et al. (2012) and Foster et al. (2015) shown in black and the Novaseq error profile as predicted by insilicoseq Lelieveld et al. (2024) shown in red. Absolute error rates for insilicoseq simulations can be found in Figure S7. Transitions are shown in blue shades and transversions in yellow shades. Chi squared values in black and red measure deviation from these two distributions respectively. We note that the expected mutational spectra comes from MA studies which use an MG1655 strain with a known pyrimidine synthesis defect repaired (*rph*+), in contrast we use the wildtype MG1655 strain which still carries this defect. Therefore the true mutational spectra may differ subtly. H) As in G but with all proportions rescaled to sum to 1 after excluding C>A mutations. The trinucleotide spectrum for DiscoSnp++, lofreq and cross-validated mutations is shown in Figure S8.

### Null Expectations of Even Positional Distribution, Mutational Spectrum and Lack of Selection

Spontaneous mutations in *E. coli* are distributed roughly evenly across the genome with only slight variation (Niccum et al. 2019), we therefore expect our observed mutations to be evenly spread rather than clustered in specific regions or positions. When using a reference-based variant calling method we find many mutations are called in multiple samples, we will call these ‘over-represented mutations’ while mutations called in only 1 sample are termed ‘other mutations’ (Figure 4). These over-represented mutations are far less likely to be called by the reference-free method (Figure 4). This discrepancy may result from our reference-free pipeline removing any SNP found on a contig which aligns to multiple genomic regions and therefore strictly avoiding SNPs which are artefacts of misalignment between homologous regions. In contrast, the reference-based pipeline used here may be more vulnerable to such errors. Further evidence that these over-represented mutations do not have a biological origin comes from their greater alternative allele depth as compared to mutations which are seen in only one sample (Figure S2). Given our simulated expectations (Figure 2) most mutations should be seen at low frequencies, in line with the mutations called in only 1 sample.

We also have prior expectations of the mutational spectrum of SNPs which should be seen in wildtype *E. coli* from mutation accumulation experiments, the current gold-standard (Lee et al. 2012; Foster et al. 2015; Long et al. 2016). A second potential model for the SNP distribution of called mutations is the error-profile of Illumina Novaseq technology used for sequencing in this study. This error-profile can be predicted by simulating reads from our genome of interest using a model trained on Novaseq reads, this was done using InSilicoSeq (Lelieveld et al. 2024). For each set of mutations we calculated the deviation from both the MA spectrum (as the mean of Long et al. (2016); Lee et al. (2012) and Foster et al. (2015)) and the Novaseq error-profile using Chi-squared statistics for which a lower statistic indicates a better fit of the data to the expected distribution.

We find that the over-represented mutations called by lofreq are a good fit to the MA spectrum (Figure 4). This is consistent with the idea that these over-represented mutations may be the result of misalignment between homologous regions which duplicated at some point prior to our experiment. If these variants are a consequence of divergence following duplication it follows that their mutational spectrum will reflect that of *E. coli* with any deviation resulting from changes in the spectrum during evolution or selection on the diverging mutations. Among the other mutations called by lofreq we find an excessive proportion of C>A mutations (Figure 4). These are the most common error under the Novaseq error-profile and so this group of mutations may be biased not by false positives from misalignment but from the calling of errors as true mutations. Calls from DiscoSnp++ are dominated by A>C mutations with neither the MA or Novaseq profiles providing a good fit to the observed spectrum (Figure 4). Once we subset for mutations which are cross-validated by both methods we find that C>A mutations are again over-represented, perhaps suggesting bias introduced by sequencing errors (Figure 4). However, after removing all C>A mutations as they are likely to be more dominated by false-positives, we see a reasonably good fit to the MA expectation (Figure 4) with the expected bias for transitions over transversions being clear (Lee et al. 2012).

When measuring mutation rates it is vital for biases not to be introduced by natural selection, this is a key aim of both MA and FA approaches though neither can reduce selection to 0 (for attempts at corrections to the effect of selection for FA see Hamon and Ycart (2012), and for MA see Wahl and Agashe (2022) *cf.* Mahilkar et al. (2022)). Given that the approach taken here, deep sequencing after a single growth cycle, allows fewer generations in which selection can take place than an MA experiment it is likely to be less biased by selection. This approach is also less likely to be biased by selection than the FA, despite both taking estimates after a single growth cycle. This is because while the FA uses phenotypically selectable markers, likely to have fitness effects (Baisa et al. 2013; Palmer et al. 2015; Harmand et al. 2017; Soley et al. 2023), deep sequencing allows all mutations to be observed most of which will impose neutral or near-neutral fitness effects (Robert et al. 2018).

Purifying selection is likely to enrich observed mutations for those with little phenotypic effect, we would therefore expect an increase in mutations in non-coding regions and, within coding regions, an enrichment for synonymous mutations when selection is acting. We can compare null expectations of mutation in the absence of selection in wildtype *E. coli* (Foster et al. 2015) to the observed mutation sets in this study to compare the influence of selection in each case. Among the mutations found in >1 sample by the reference-based lofreq pipeline (‘Lofreq - Over-represented’) we find a strong signature of selection with strong enrichment for non-coding mutations and, within coding regions, for synonymous mutations (Figure 5). This is consistent with the idea that these ‘mutations’ are the result of misalignment between diverged, homologous regions of the genome as following the duplication of a region any mutations accumulating will be subject to selection and will therefore show the observed enrichment patterns. In contrast, the mutations found in only 1 sample by lofreq (‘Lofreq - Other’) are a much better fit to the null expectation in the absence of selection (Figure 5).

**Figure 5:**
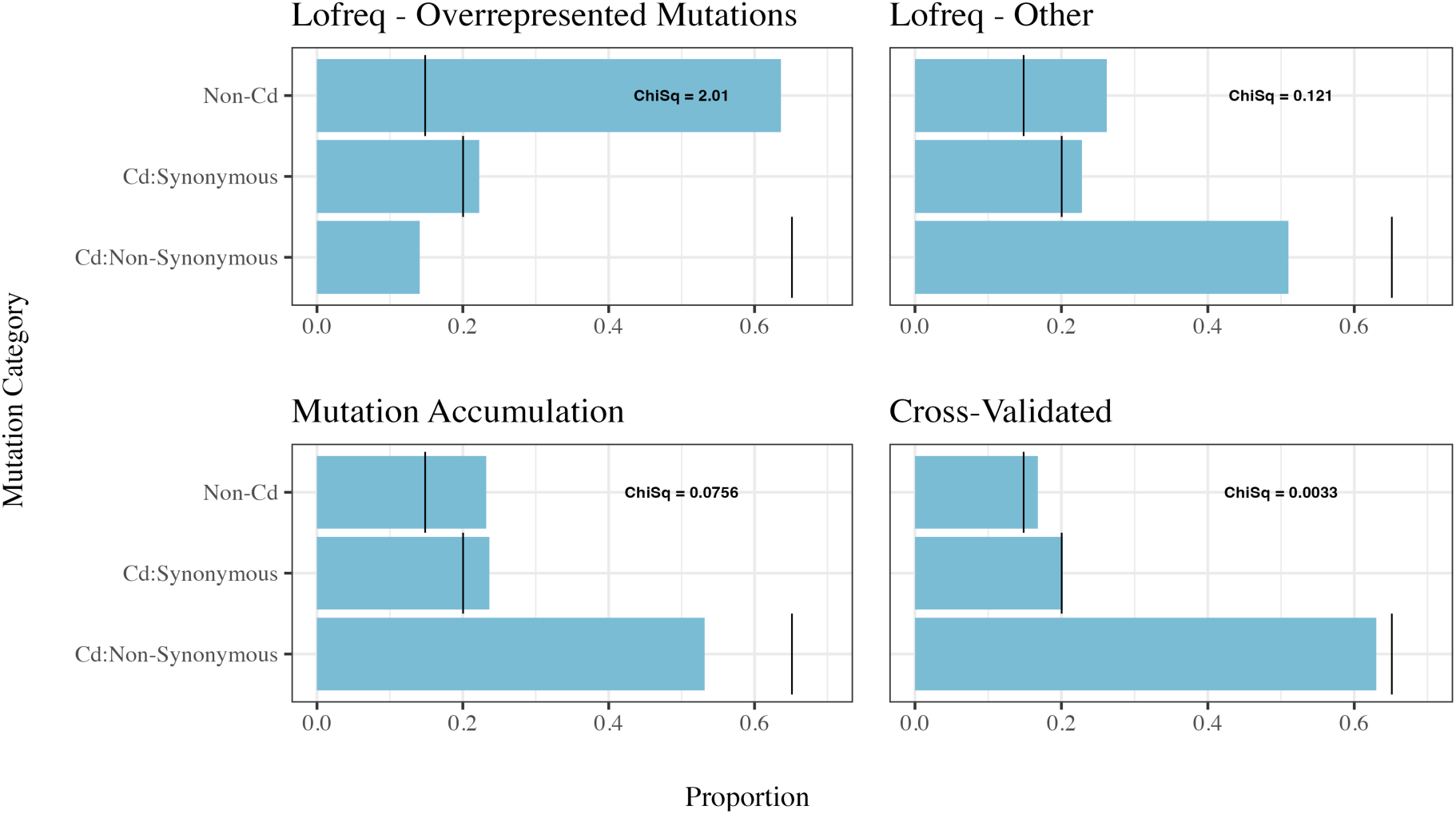
Cross-Validated mutations fit well to null expectations in the absence of selection. The proportions of SNP mutations classified as ‘non-coding/intergenic’, ‘coding: synonymous’ and ‘coding: nonsynonymous/nonsense’ are shown in each sub plot. Clockwise from top left the mutations shown in each plot are those mutations called by lofreq in more than 1 sample (Overrepresented) (total N = 6110), lofreq mutations called in only 1 sample (Other) (total N = 504), mutations cross-validated by being called by lofreq and DiscoSnp++ (total N = 119) and mutation accumulation results from Foster et al. (2015) (total N = 246). In each subplot black bars show the null expectations of mutation distribution among these 3 categories if mutations are randomly distributed in the genome, these predictions are taken from Foster et al. (2015). Chi-Squared values measure how well the observed distribution fits this null expectation with lower values indicating a better fit.

Despite this, some signature of selection remains, indicating either that false-positives from diverged, homologous regions are also found in this mutation set or that substantial purifying selection is acting on true mutations occurring during the single growth cycle in this study. Given that the distribution of mutations from MA experiments with wildtype *E. coli* MG1655 *rph*+ (Figure 5) are likely to experience more selection on true mutations, yet fit better to the null expectation, we believe the deviations seen in ‘Lofreq - Other’ are more likely to result from from the misalignment of diverged, homologous regions than selection during the experiment. Mutations cross-validated by both the reference-based lofreq and reference-free DiscoSnp++ pipelines are a very close fit to the null expectations of no selection (Figure 5) suggesting that selection is minimized by this method. However, sequencing errors will also be subject to no selection and therefore bias introduced by sequencing error could also produce a good fit to our null expectations.

While testing for the influence of purifying selection, it cannot distinguish the influence of false-positives from sequencing errors. In contrast, comparisons to the expected mutational spectrum (Figure 4) can distinguish the influence of sequencing error, as this profile differs from the expected mutational spectrum of *E. coli*. Alternative allele depth is also likely to be elevated in false-positives from misalignments (Figure S2). Applying these three tests is therefore an effective way to test for the influence of these two key false-positive sources. From the mutational spectrum of cross-validated mutations we identified a disproportionately high fraction of C>A mutations (Figure 4). We therefore compared variant allele depth and signatures of purifying selection on these C>A mutations and all other mutations separately, finding that neither C>A mutations nor all non-C>A mutations had a strong signature of purifying selection and do not vary in variant allele depth (Figure S3). This may suggest that these excess C>A mutations derive from sequencing-errors rather than misalignment. Another potential source of false-positive C>A mutations is oxidative damage to nucleotides during DNA extraction (Stoler and Nekrutenko 2021) when followed by sequencing by synthesis. This can result in C>A mutation calls as G nucleotides are most vulnerable to oxidative damage and, once oxidised, readily pair with A nucleotides resulting in GC>TA mutation calls (Maki and Sekiguchi 1992). Alternatively, it is possible that these C>A mutations are not false positives and instead reflect a true bias towards C>A mutations under the conditions of this experiment, given the strength of this bias we believe this alternative explanation is unlikely to be true.

### Genome-Wide Evidence of Density Associated Mutation Rate Plasticity

Having subset our data to reduce bias introduced by false-positive variant calls we can now test the relationship between mutation rate and population density in wildtype *E. coli*. While a negative association between mutation rates and population density has been observed at various marker loci using a FA approach, truly genome-wide data has yet been reported for this phenomenon (Krašovec et al. 2014; Krašovec et al. 2017; Krašovec et al. 2019). To test for the presence of density associated mutation rate plasticity (DAMP) we fitted a Poisson model describing the number of mutations per sample as a function of the log_2_ transformed population density (Figure 6). When filtering out C>A mutations from our cross-validated dataset we find a strong DAMP slope of −0.72 (z = −3.3, *P* = 0.0011, Figure 6A). In line with our hypothesis that the excessive C>A mutations derive from non-biological processes such as sequencing errors or oxidative damage during DNA extraction we find a weaker DAMP slope of only −0.3 among the cross-validated C>A mutations (Figure 6E). However, it is also worth noting that C>A mutations are less responsive to density than other mutational types (e.g. A>G transitions) and so may show a weaker slope even if they are true-positives (Gifford et al. 2024; Green et al. 2025). Our observation of much weaker, or even reversed, DAMP among the raw mutations called by both the reference-based lofreq pipeline and the reference-free DiscoSnp++ pipeline validate the use of cross validation between these pipelines in order to enrich for true-positive mutations in order to observe biologically relevant trends.

**Figure 6:**
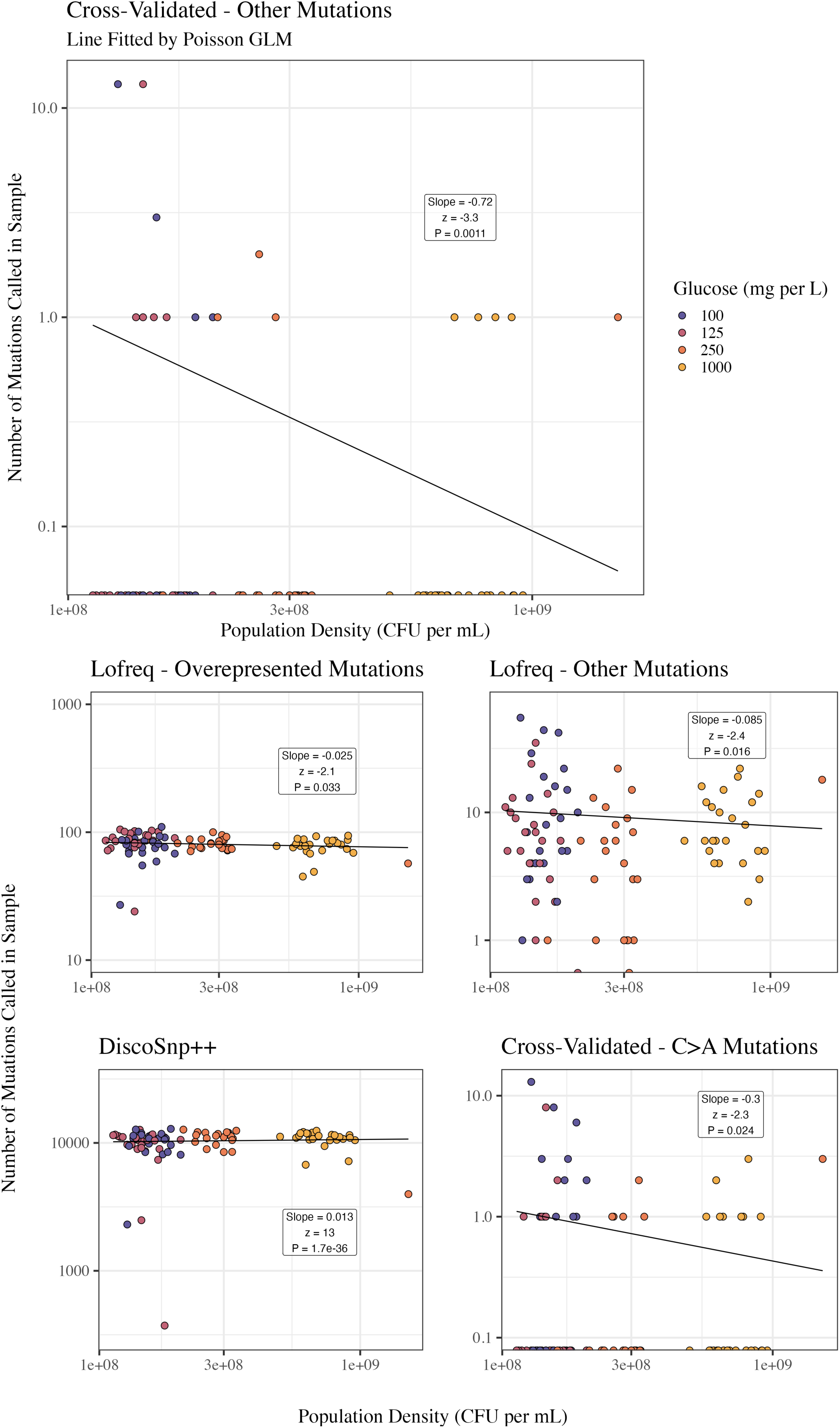
Cross-Validated, non-C>A, mutations show the strongest association between mutation count and population density. For each set of mutations a glm model using Poisson family with a log link is fitted describing number of mutations per sample as a function of the log_2_ transformed population density. Slope, z values and associated p values comparing the slope to 0 are shown for each fitted model. Note log scaled x and y axes on all subplots. All y-axes cover a roughly 100-fold range to make slope magnitude visually comparable.

The number of observed mutations per sample will be biased by the sequencing coverage of that sample. If coverage is negatively correlated with population density then the observation of DAMP in this data may be an artifact. To test this alternative hypothesis we measured the correlation between population density and sequencing coverage. We find a shallow negative relationship between population density and sequenced bases per sample (Figure S5). To test the extent to which this variation in sequencing coverage may affect the observations in Figure 6 we simulated a constant mutation rate with sequencing coverage following the trend in Figure S5A. These simulations take the same form as those shown in Figure 2. These simulations predict no significant relationship between observed mutation counts and population density (slope = −0.031 ± 0.14), z = −0.45, *P* = 0.66, Figure S4). It is therefore unlikely that the slopes seen in Figure 6 are the result of variations in sequencing coverage.

## Discussion

This study highlights the great promise of modern sequencing technology to overcome the pitfalls of current mutation rate assays. We also outline potential biases of a deep sequencing based approach and propose methods for identifying and minimising bias. Using simple calculations to estimate expected mutant counts across a variety of parameters we quantify the minimum necessary sequencing depth required for such approaches to be ∼1000-fold coverage (Figure 2). We sequence a set of *E. coli* samples to this depth finding that while sequencing and alignment errors are frequent these can be minimised by cross-validating mutations called by diverse methods (Figure 4, Figure 5). These identified mutations are then used to present the first evidence of a genome wide signature of density associated mutation rate plasticity (Figure 6).

A key challenge here is overcoming biases introduced by sequencing errors. It would be possible to reduce this noise by using paired end sequencing with small insert size so that mutations can be verified on both the forward and reverse reads (Chen-Harris et al. 2013; Fusco et al. 2016; Preston et al. 2016). Progress will also be made in this respect as error rates of next generation sequencing technology continue to fall. It would also be beneficial to increase sequencing coverage to ≥10,000 fold coverage in order for a greater number of mutants to be observed. Strong positive-controls could also be implemented both by spiking in known mutant populations at known frequencies prior to sequencing and by computationally ‘spiking’ known mutations into the raw or trimmed reads. Finally, as we proposed that the over-abundance of C>A mutations among the cross-validated group may be the result of oxidative damage during DNA extraction it would be useful to test alternative extraction methods to minimise this source of noise.

When assessing environmental effects on mutation rates it is important to hold all other demographic variables constant. Population density in this study was controlled by provisioning populations of the same volume with differing glucose concentrations and thereby facilitating a greater or lesser number of generations. In contrast if total volume were varied so that glucose concentration could be varied without varying glucose mass then all populations could be allowed to reach differing densities while the number of generations and hence final population size were held constant. This may help to avoid biases both in the effect of selection and in differential sequencing depth by treatment group (Figure S5).

We hope that this study will motivate further work into leveraging the potentials of deep sequencing to modernise microbial mutation rate assays. Common methods for mutation rate measurement have not been significantly modified in many decades and experience a trade of between genomic scale (1 target vs whole genome) and experiment run time while also being biased by selection (Table 1). Here we have outlined how deep sequencing has the potential to combine the genome wide scale of MA assays with the short run time of FA assays while also reducing selection bias.

## Methods

### Experimental Methods

Experimental design is similar to a fluctuation assay as described in (Krašovec et al. 2019). A single ice scrape of MG1655 wildtype *E. coli* was revived in 5mL LB until turbid. This was then diluted 100,000 fold into 4 overnight cultures of DM media supplemented with 100, 125, 250 & 1000 mg/L glucose respectively. These overnight cultures were then diluted to a population size of 6500, 4500, 3500 & 1000 respectively for the 4 glucose concentrations. (The negative correlation between glucose concentration and initial population size was unintentional and is perhaps an inconvenient confounding variable). 24 cultures of 10mL were incubated for 25 hours in each glucose concentration. From these cultures 700**μ**L was frozen at −80, 750**μ**L was plated on rifampicin to test the mutation rate to rifampicin resistance, 100**μ**L was diluted by a factor of 10^5^ before plating on non-selective agar to assess the final population size and 8.3mL was used for DNA extraction using a ‘Macherey-Nagel™ NucleoSpin™ Microbial DNA kit for DNA from microorganisms’. This process is outlined in Figure 7.

**Figure 7:**
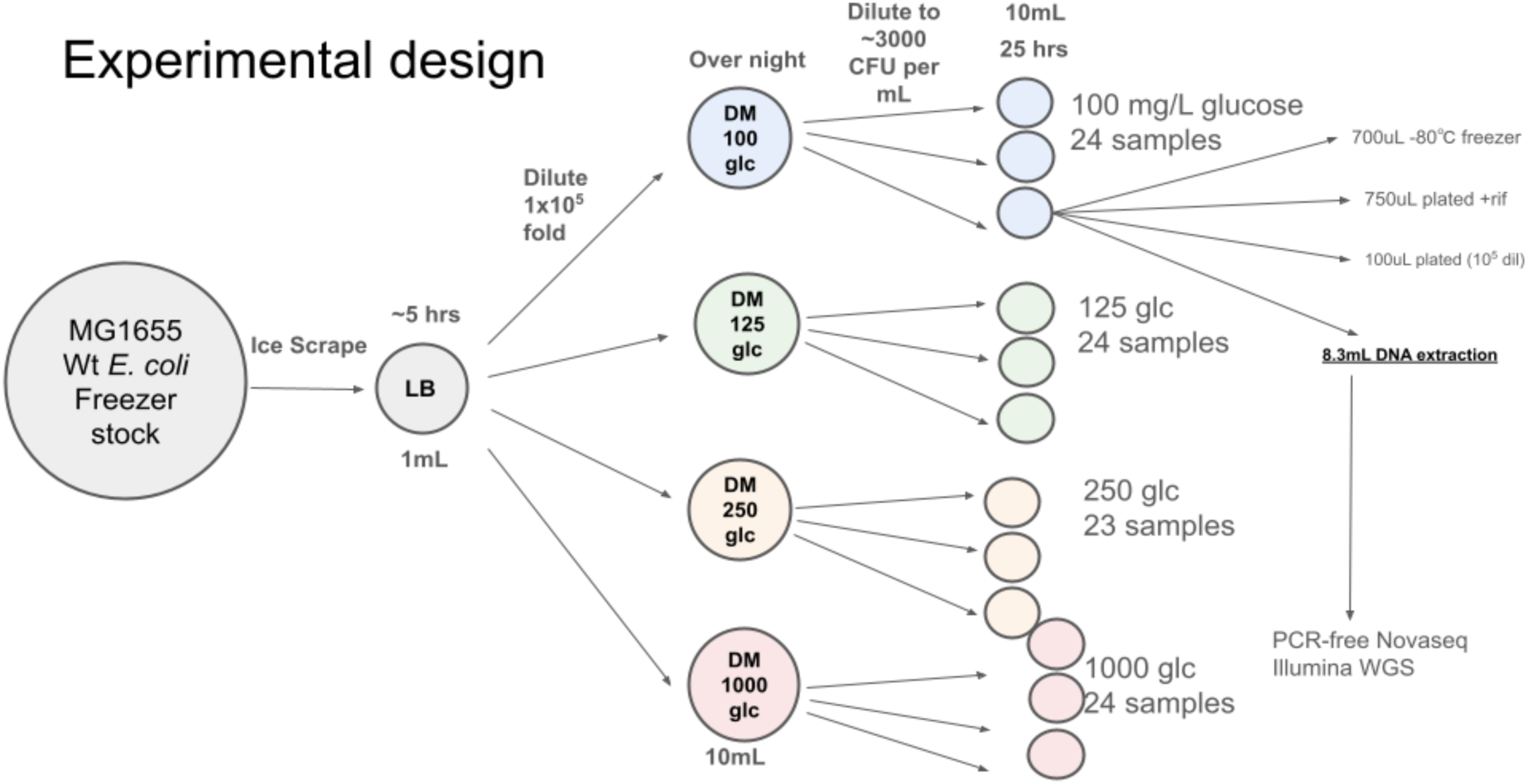
Illustration of experimental design.

### Illumina Novaseq PCR-free sequencing

The libraries for this project were constructed by the Technical Genomics Team at the Earlham Institute, Norwich, UK using the KAPA High Throughout Library Prep Kit (Roche Part No: KK8234/07961901001). Where possible 1µg of genomic DNA was sheared to 450bp using the Covaris ML230 Sonicator (Covaris) and the ends of the DNA were repaired; 3’ to 5’ exonuclease activity removed the 3’ overhangs and the polymerase activity filled in the 5’ overhangs creating blunt ends. A single ‘A’ nucleotide was added to the 3’ ends of the blunt fragments to allow for the ligation of barcoded adapters (6bp - Perkin Elmer NEXTFLEX DNA Barcodes 1-48 (NOVA-514101/2/3/4)) or (12bp - Perkin Elmer NEXTFLEX-HT (NOVA-51474/5/6/7)) at a concentration of 6µM prior to a 0.8x clean up using Beckman Coulter AMPure XP beads (A63882). The size of the libraries was estimated using an Agilent High Sensitivity DNA chip (5067-4626) and the concentrations were quantified by fluorescence with a High Sensitivity Qubit assay (ThermoFisher Q32854).

### Variant Calling Pipelines

Both the reference-free and reference-based pipelines are shown in Figure 3. All raw reads were initially trimmed and adapter content removed using trimmomatic (Bolger et al. 2014). Due to uneven base frequencies within the first 10 basepairs of each sample, identified by fastqc (https://www.bioinformatics.babraham.ac.uk/projects/fastqc/), we applied a headcrop of 10 to all reads. Trimmomatic was run as follows:

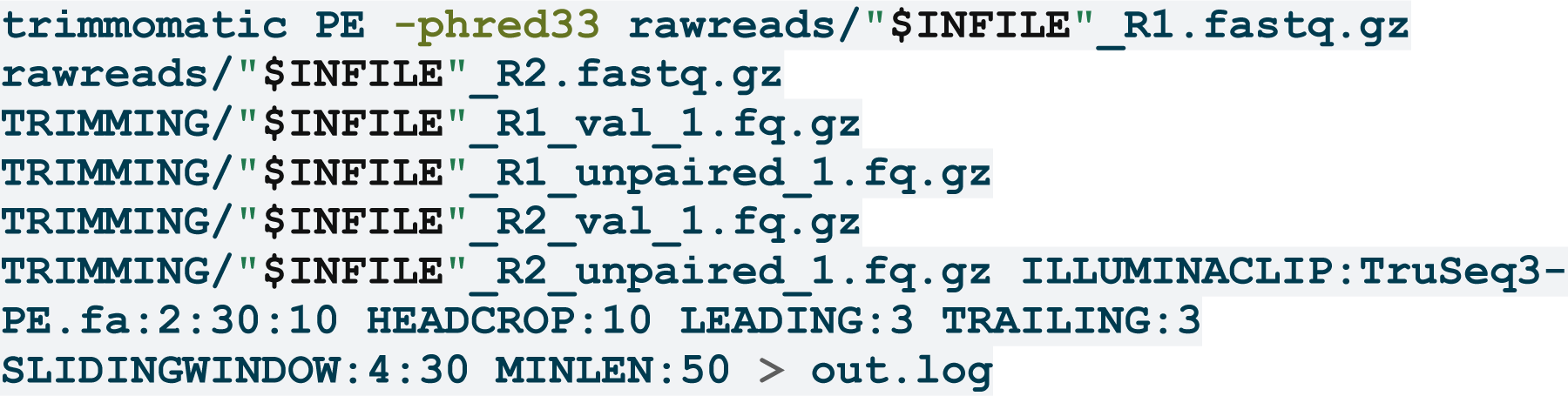

Our reference-free approach used DiscoSnp++ (Peterlongo et al. 2017). DiscoSnp++ uses de Bruijn graphs to identify SNPs as bubbles in a reference free manner. We allowed for contig extension around these SNPs; after variant calling & contig extension contigs are aligned to the reference genome using BWA-mem. To avoid the calling of mutations where multiple homologous regions may be aligned to one another we discarded all contigs with >1 alignment locus. Mutations were only called where ≥2 reads supported both the reference and alternative allele. Finally mutations were annotated using gdtools from breseq (Deatherage and Barrick 2014). This pipeline is shown here:

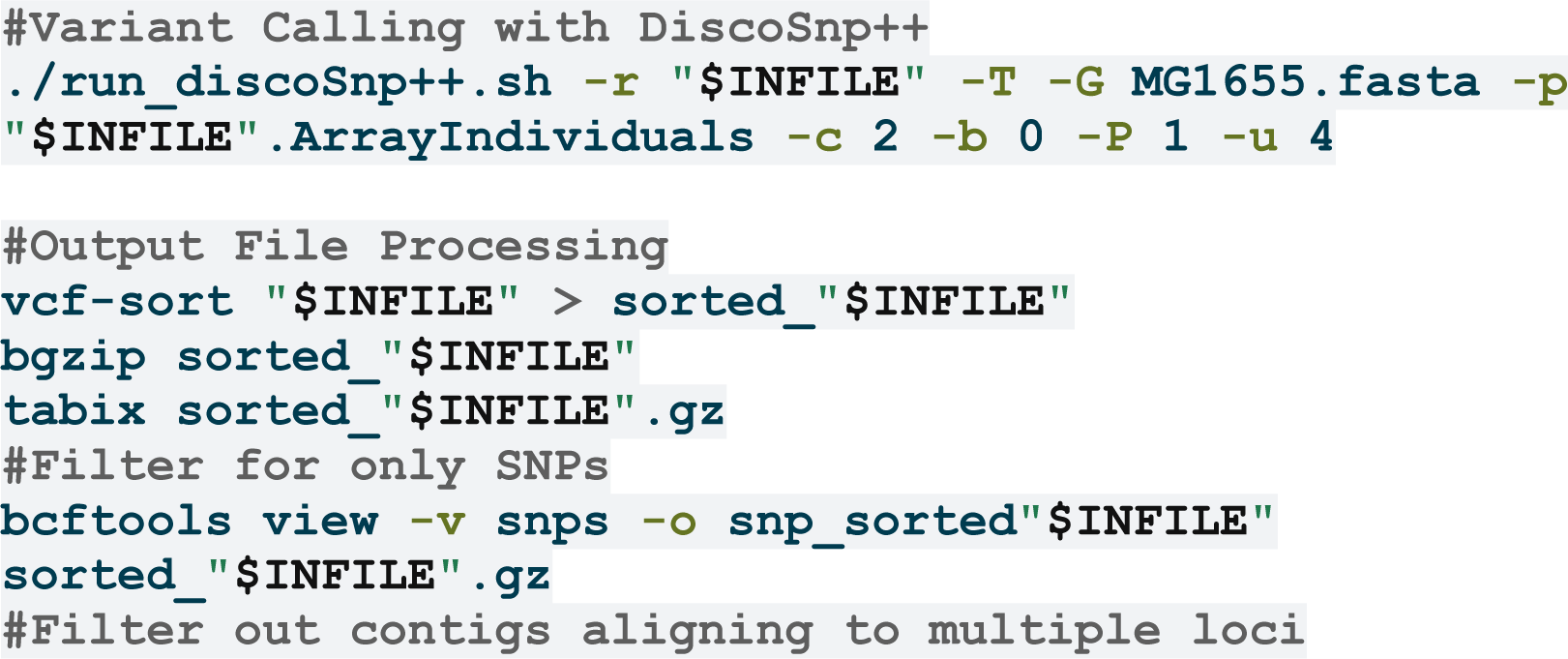
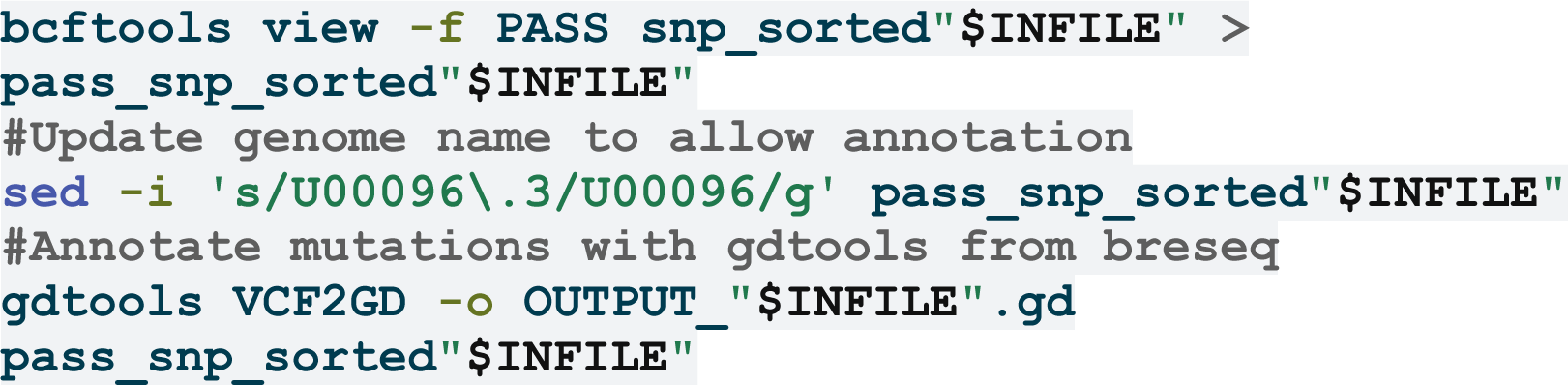

Our reference-based pipeline was based on that used by (Jago et al. 2025) with trimmomatic used for trimming (see above) as opposed to trimgalore. This pipeline calls variants using both mpileup (Danecek et al. 2021) and lofreq, a tool specialised for calling rare variants (Wilm et al. 2012). Mutations called by this pipeline were filtered for only those with ≥1 reads in both the forward and reverse direction supporting the reference and alternative alleles. This pipeline is as follows and can also be found in the original publication by (Jago et al. 2025):

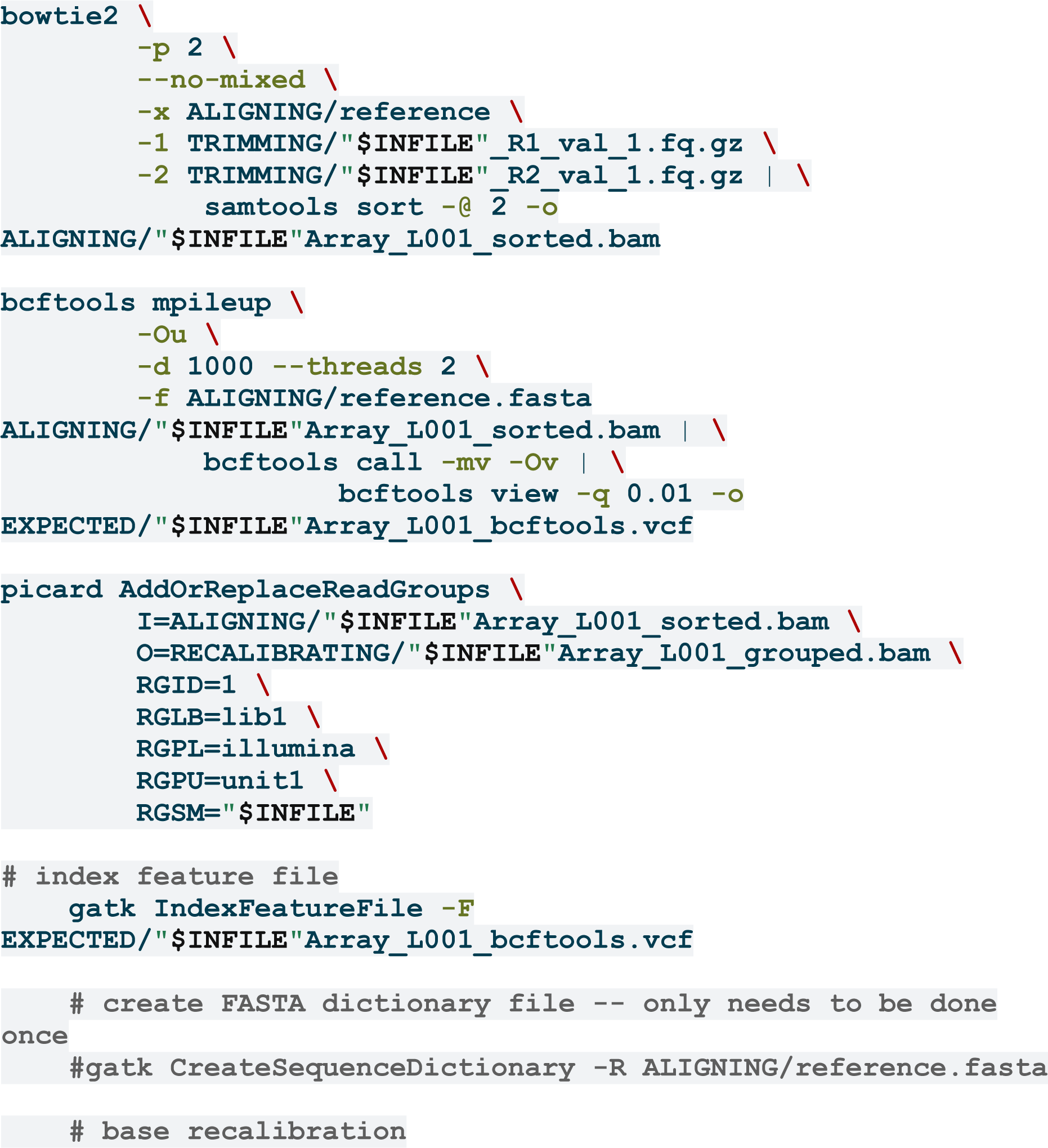
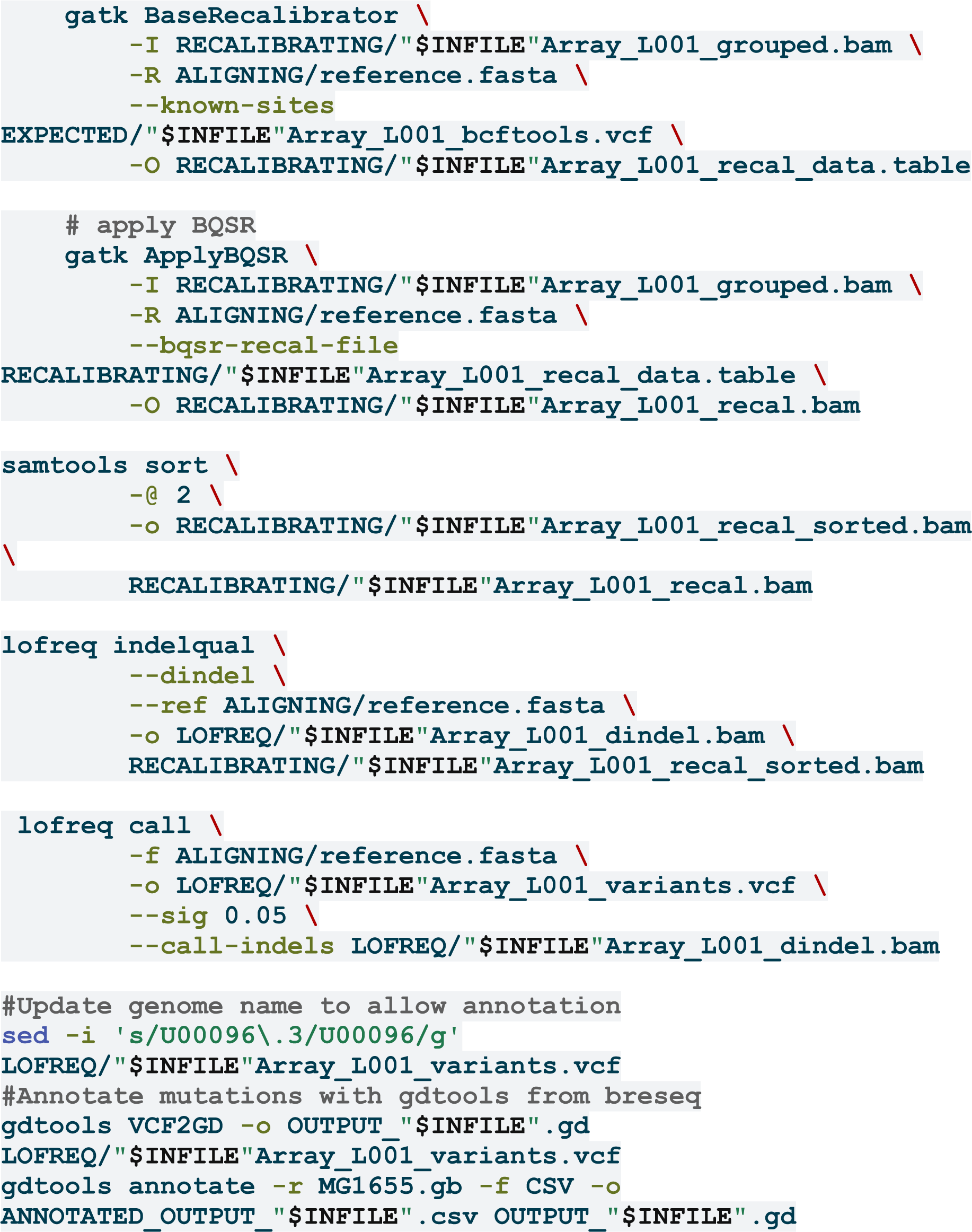

### Simulated Reads

1 million Novaseq reads were simulated using InSilicoSeq (Lelieveld et al. 2024) using the following options:

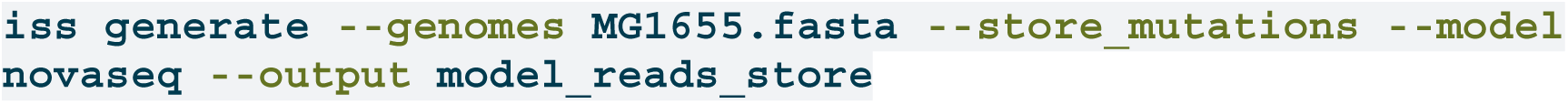

### Analysis and Visualisation

All analysis and visualisation was performed in R version 4.3.1 (R Core Team 2023) using packages emdbook (Bolker 2023), tidyverse (Wickham et al. 2019), magrittr (Bache and Wickham 2022), plyr (Wickham 2011), vcfR (Knaus and Grunwald 2017), ggVennDiagram (Gao and Dusa 2024), readxl (Wickham and Bryan 2023), ggbeeswarm (Clarke et al. 2023), cowplot (Wilke 2024), gridExtra (Auguie 2017), magick (Ooms 2024), DiagrammeR (Iannone 2016; Iannone and Roy 2024), rsvg (Ooms 2023), svglite (Wickham et al. 2023), png (Urbanek 2022), ggpubr (Kassambara 2023) and PNWColors (Lawlor 2020).

## Acknowledgements

The authors would like to thank Matthew Thomas, Asher Leeks and Penny Kahn for insightful discussions on this manuscript. The authors would also like to thank the Computational Shared Facility at The University of Manchester and the Advanced Research Computing facility ‘Sockeye’ at The University of British Columbia (UBC Advanced Research Computing 2019).

## Supplementary Figures

**Figure S1:**
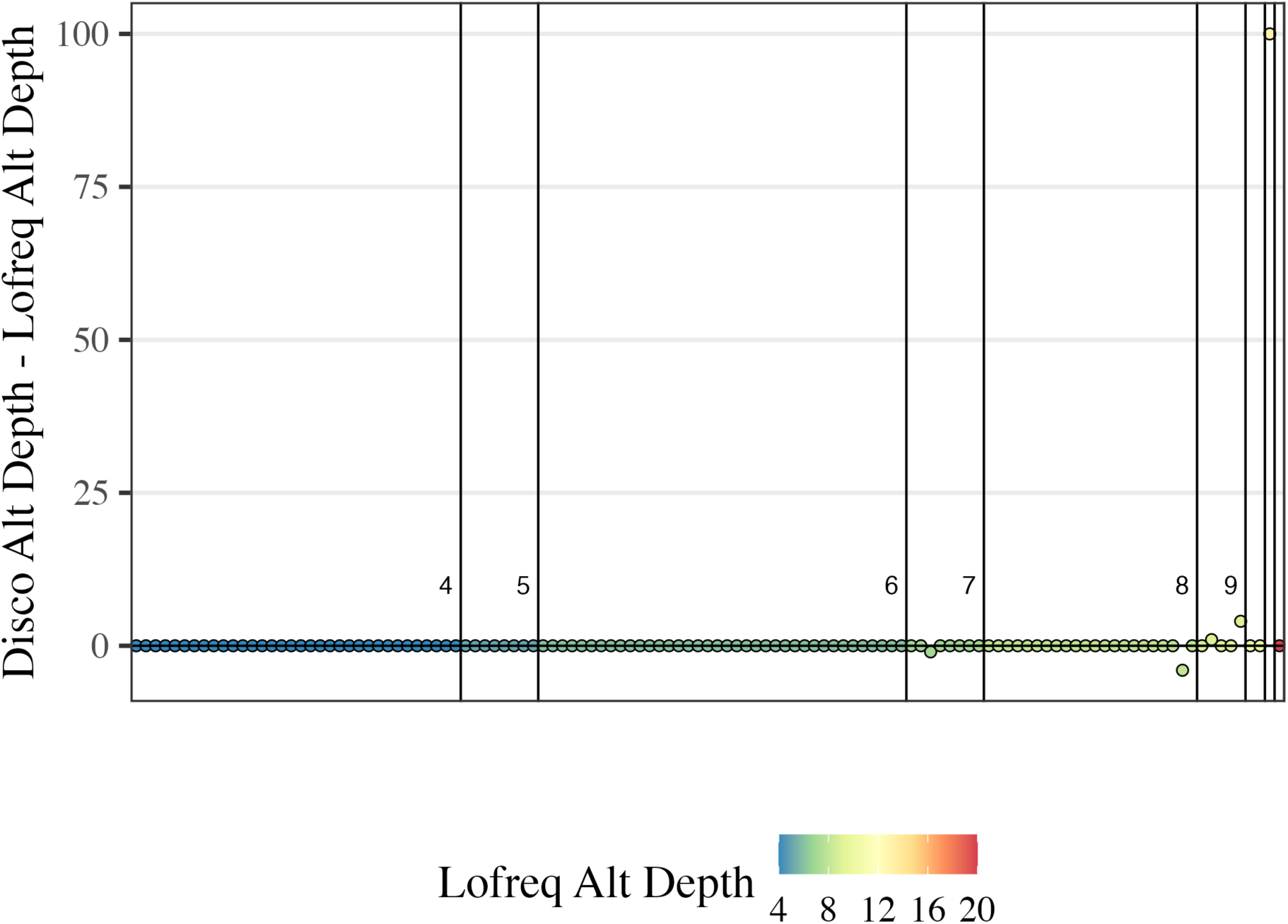
Depth of variants called by both lofreq and discosnp++ are generally in agreement. Each point represents a single mutation with the difference between alternative allele depth predicted by DiscoSnp++ and lofreq shown on the y-axis. Points are coloured by the alternative allele depth predicted by lofreq. Vertical lines separate each group of mutations with the same lofreq depth. For each group with >2 mutations the lofreq depth is shown above the relevant mutation points.

**Figure S2:**
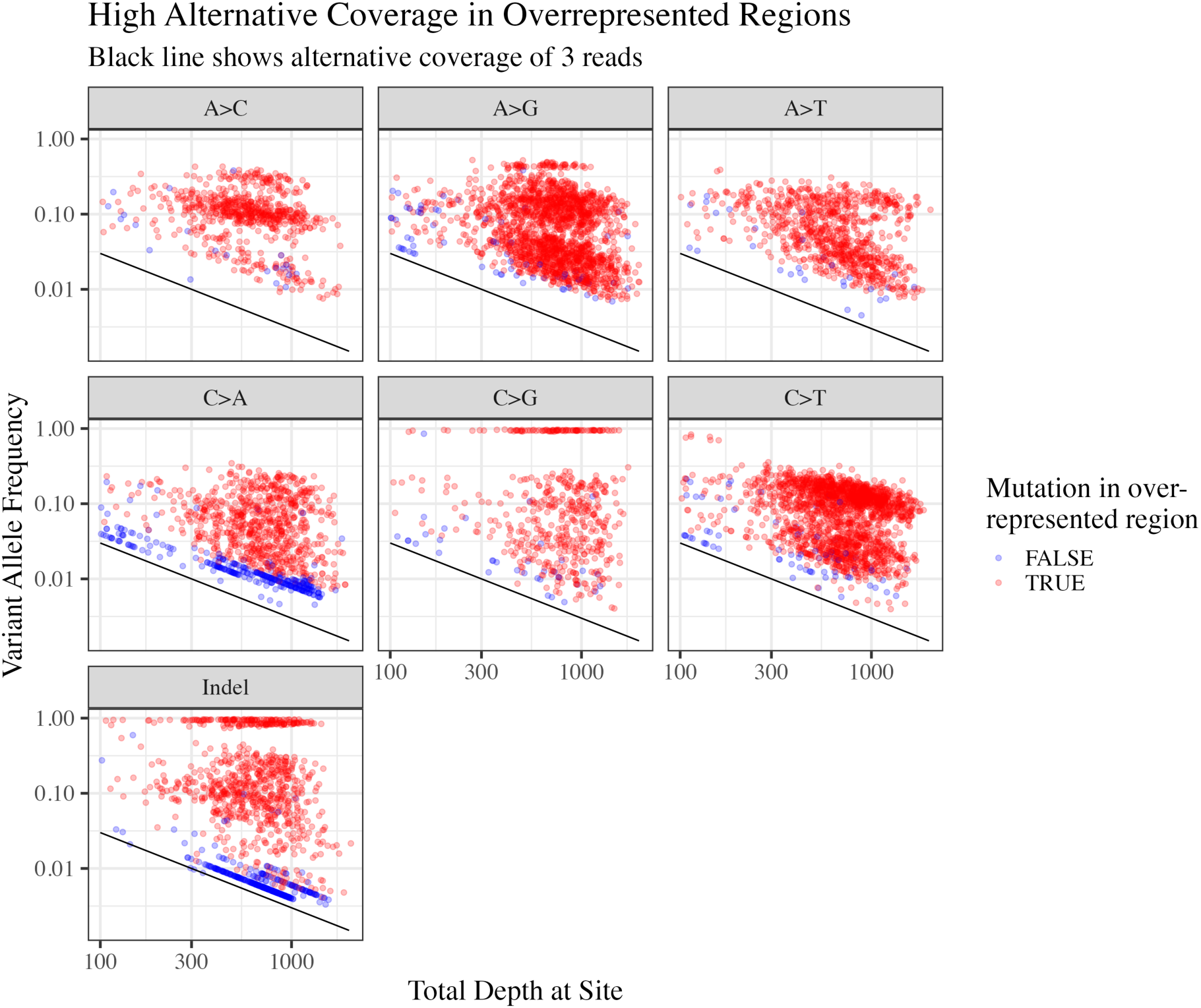
Over-represented mutations are seen at greater depth than mutations called in only one sample. For each mutational class mutations called by the reference-based pipeline are shown with total depth (coverage) on the x axis and variant allele frequency on the y axis. Mutations with the same variant allele depth will form diagonal lines, an example line for variants supported by 3 reads is shown by the diagonal black lines. Mutations called in more than 1 sample (over-represented) are shown in red and those called in only a single sample are shown in blue.

**Figure S3:**
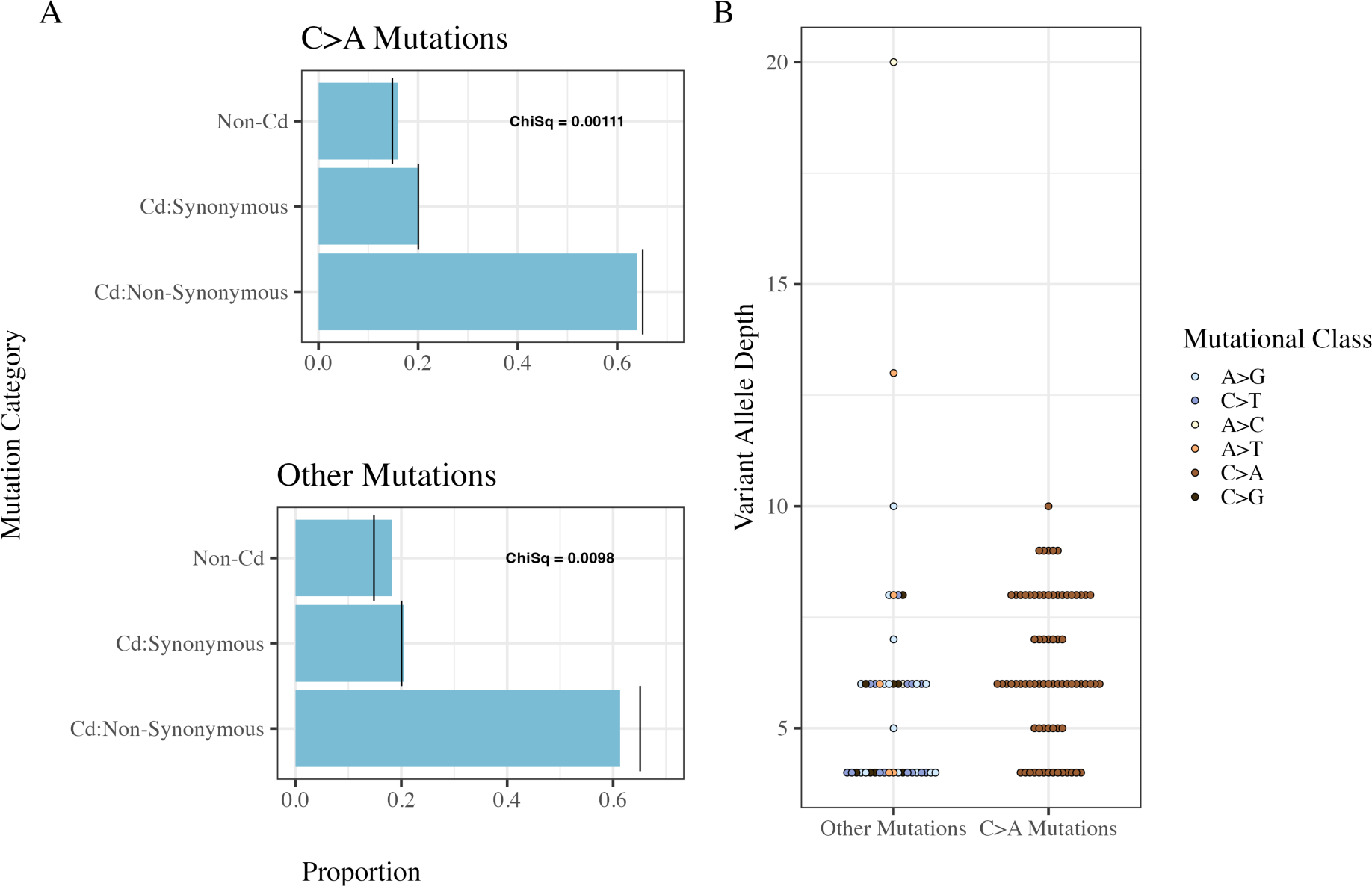
Cross-validated C>A mutations do not differ from other mutational classes in signatures of purifying selection (A) or variant allele depth (B). Chi-Squared values in panel A compare the observed proportions of mutational cateogries to the expected proportions as given in Foster et al. (2015) with lower values indicating better fit between these distributions. In panel B there is no significant difference in means between variant allele depth of C>A and other mutational classes (t=1.3, *P* = 0.21).

**Figure S4:**
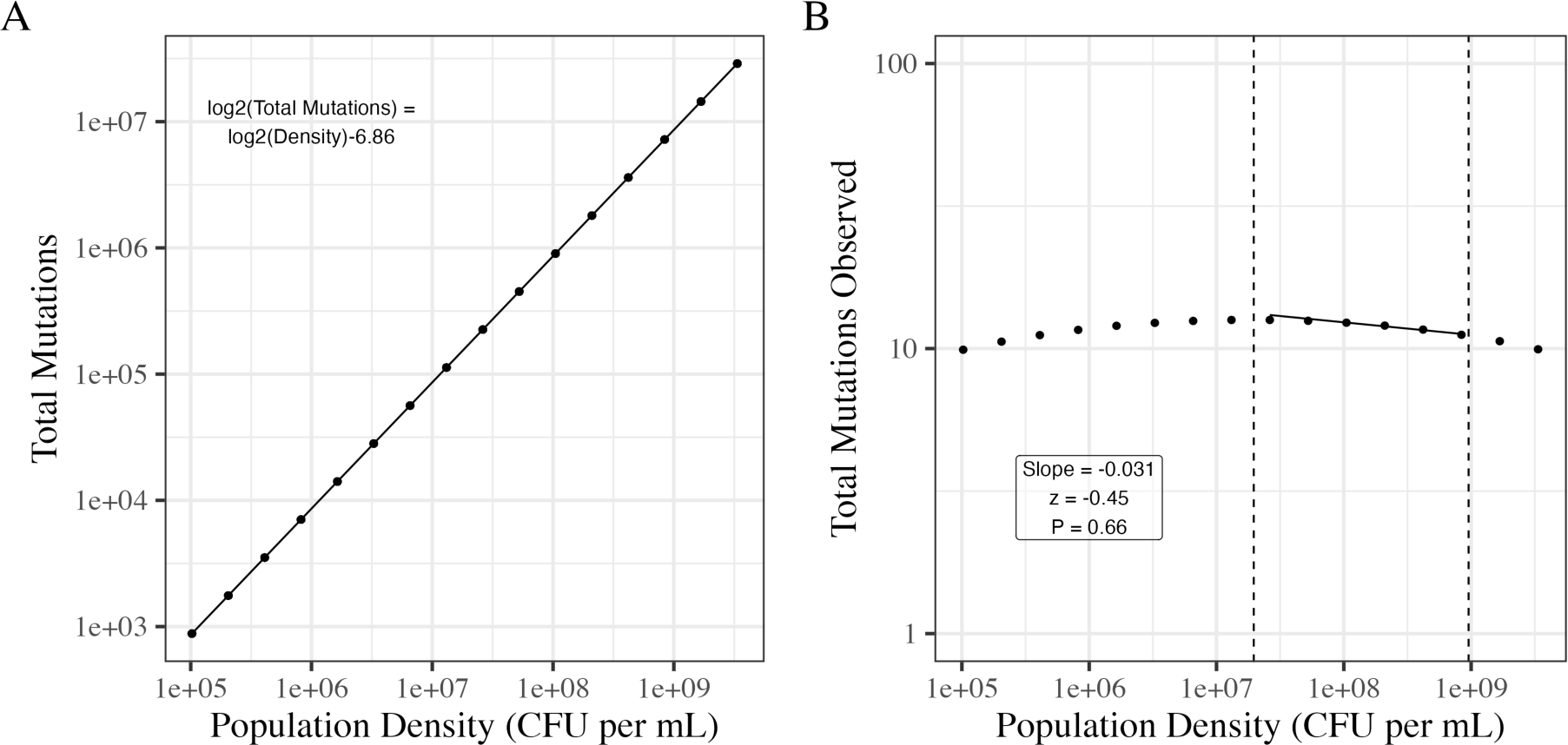
Null expectations of DAMP calculated accounting for variable sequencing coverage. Simulations are the same as those shown in Figure 2 with the modification that sequencing coverage is allowed to covary with population density following the negative line of best fit shown in Figure S5A. All simulations vary from 10 to 25 generations, are initiated by 1000 initial cells and are assumed to take place in 10mL, reflecting experimental data from this study. Therefore population densities shown here are the simulated population size divided by 10mL. A) There is a log-log linear relationship between the population density and the expected total number of mutations occurring in the population. B) We initially see a slight increase in mutant observations as density increases as a result of the positive relationship seen in panel A. However, at higher density the effect of decreased coverage reduces the number of mutant observations. A Poisson model is fitted describing the number of observed mutations as a function of the log2 transformed population density, as in Figure 6, to data within the range of population densities observed in our experimental data (marked by vertical dashed lines). Observed mutation counts are estimated as the total expected number of mutant clones observed at depths of 1 to 100. These simulations assume ∼1000x coverage therefore 100 variant reads implies a 10% mutant frequency for an individual clone, far higher than should be possibly achieved in this experiment.

**Figure S5:**
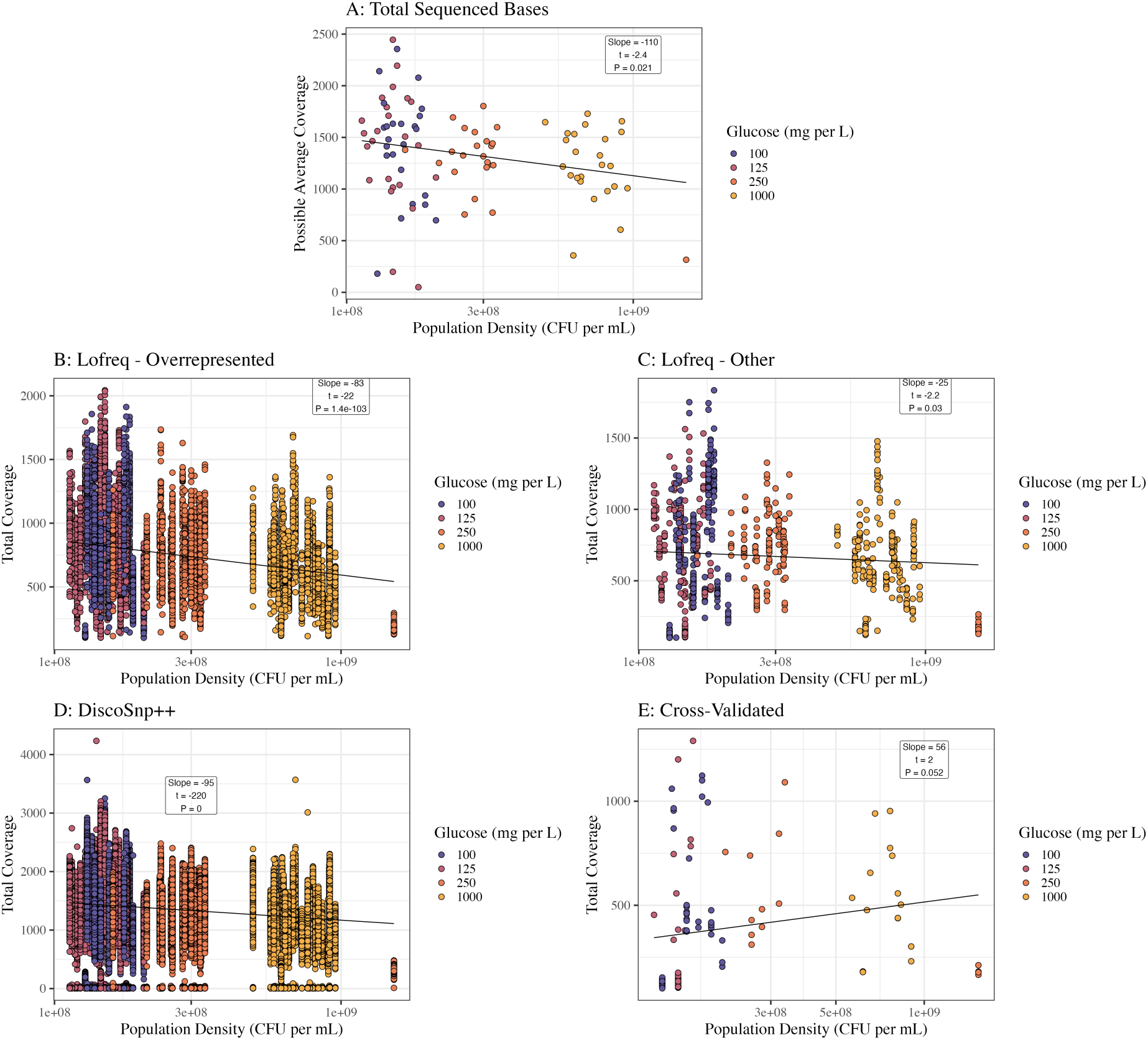
Sequencing coverage (reads per base) as a function of population density (all slopes with density log2 transformed). In panel A possible average coverage per sample is calculated as total sequenced bases divided by the genome size of MG1655 (4.6e6). In panel E total coverage is taken from lofreq estimates though note that these are generally identical to discoSnp++ estimates (Figure S1). In panel A each point represents a single sample whereas in panels B-E each point represents a single mutation with total coverage being the reference allele depth + the variant allele depth.

**Figure S6:**
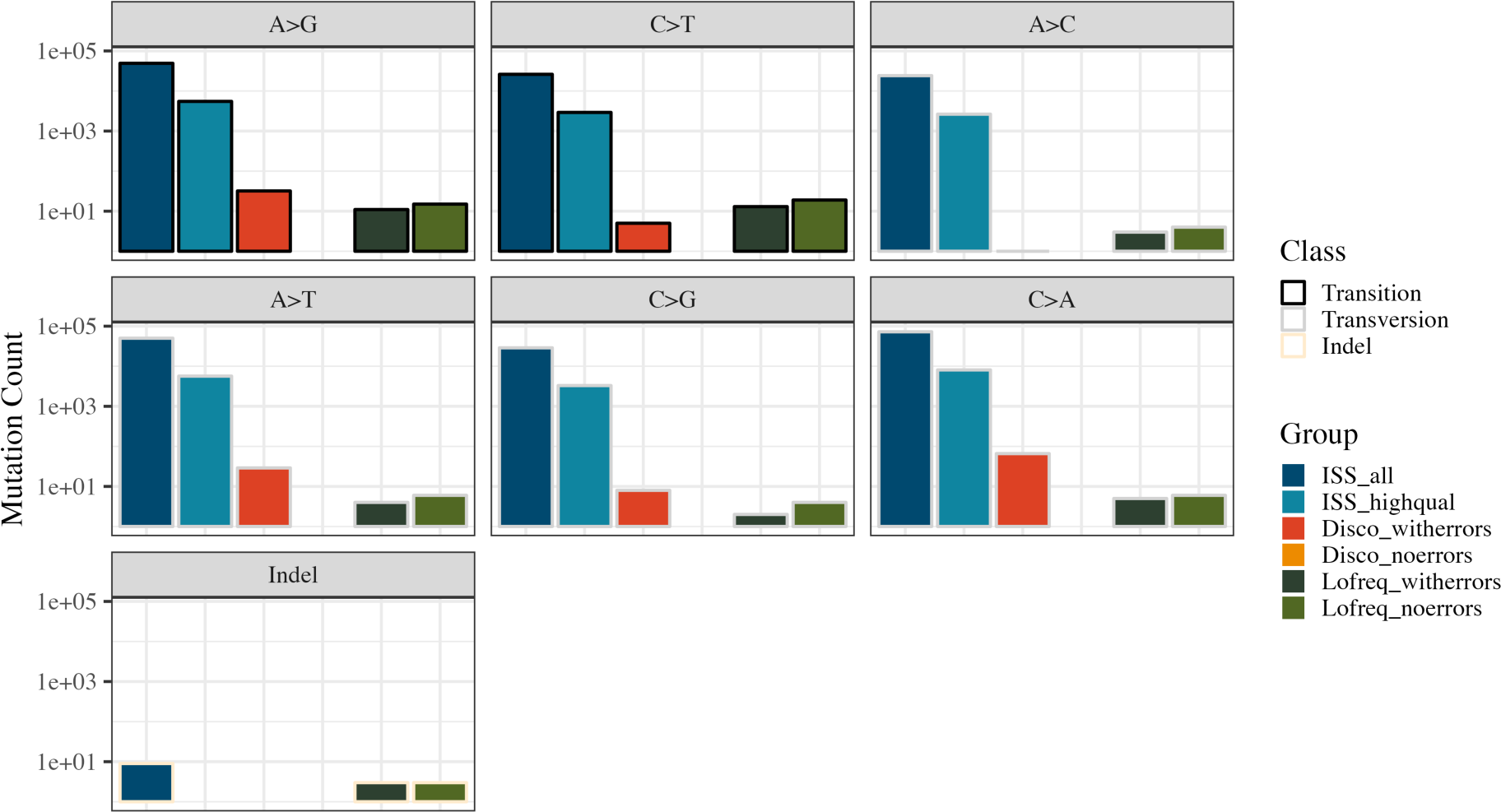
Simulated novaseq reads from InSilicoSeq. Each bar shows the count of mutations in a given category, note log scale y-axis. Variant calls form the simulated reads were performed after trimming as with all read data. From left to right the bars are: Dark blue - all simulated errors on 1 million simulated reads (total N = 250894); Light blue - simulated errors with quality >30 (total N = 28131); Dark orange - mutations called by DiscoSnp++ from InSilicoSeq reads (total N = 141); Light orange - mutations called by DiscoSnp++ from InSilicoSeq reads with all errors manually corrected (total N = 0); Dark green - mutations called by lofreq from InSilicoSeq reads (total N = 41); Light green - mutations called by lofreq from InSilicoSeq reads with all errors manually corrected (total N = 57). 1 million reads were simulated providing ∼30X coverage of the *E. coli* genome.

**Figure S7:**
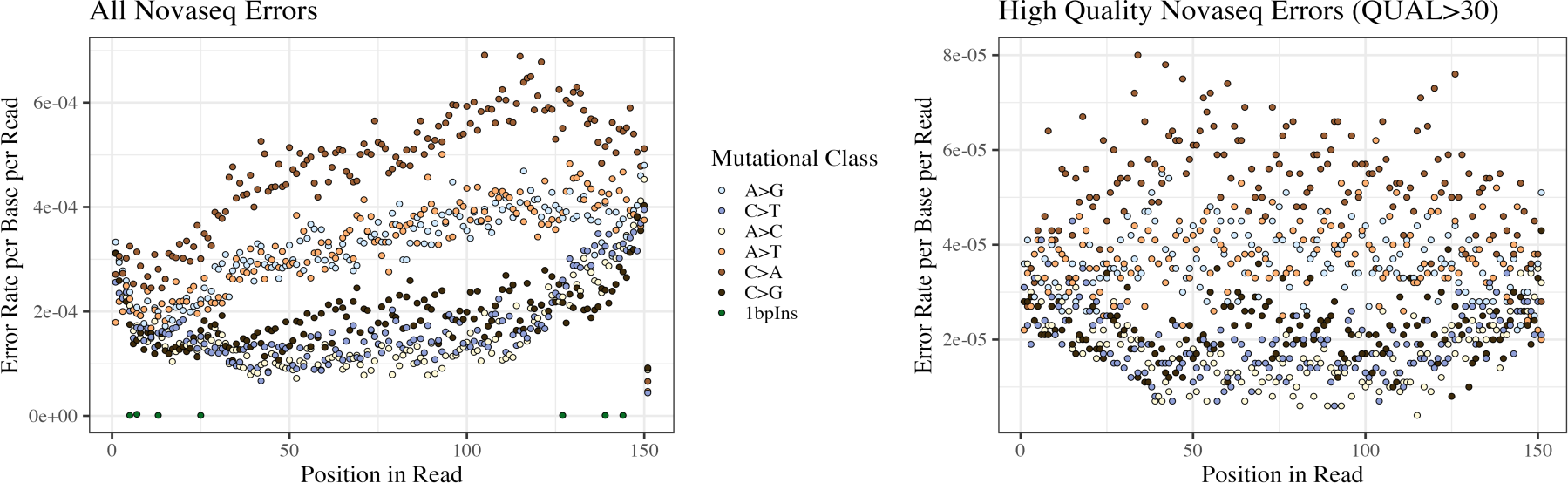
Error rates of simulated Novaseq reads by InSilicoSeq. Error rates per base per reads are calculated for all simulated errors (left) and for those errors with quality score > 30 (right). Error rates are shown for each SNP class separately.

**Figure S8:**
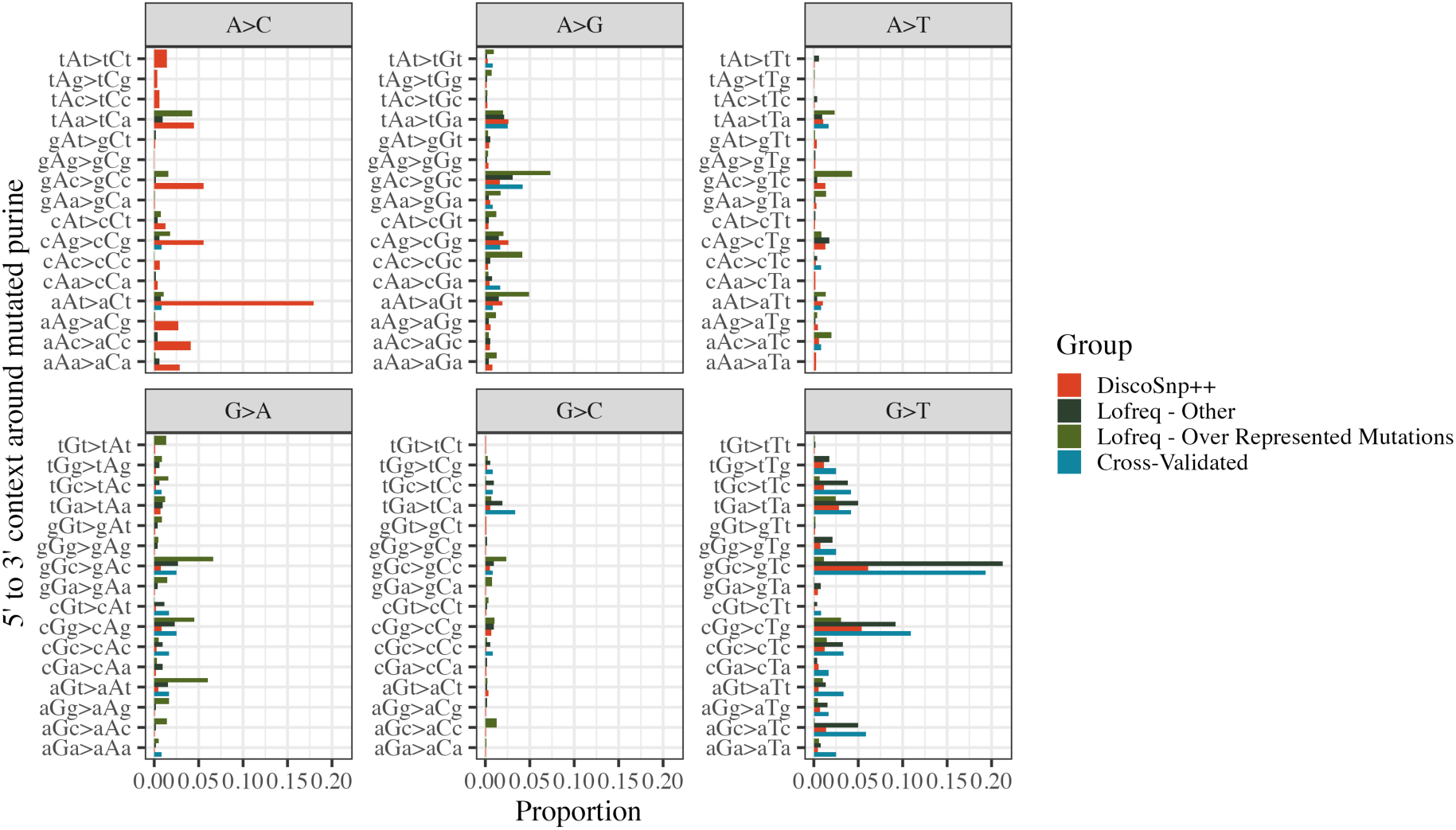
Trinucleotide context around the focal mutated purine as called by each pipeline. Note the renaming of C>X mutations as G>X so as to focus on the mutated purine. Bias within each mutational class is evident.

## Notes

https://osf.io/gztjk/

### Competing Interest Statement

The authors have declared no competing interest.

### Summary of Updates

A link to analysis code and data on the OSF has been added

### Funding

- UK Research and Innovation: MR/T021225/1

